# Astrocytic RNA editing regulates the host immune response to alpha-synuclein

**DOI:** 10.1101/2024.02.26.582055

**Authors:** Karishma D’Sa, Minee L. Choi, Aaron Z. Wagen, Núria Setó-Salvia, Olga Kopach, James R. Evans, Margarida Rodrigues, Patricia Lopez-Garcia, Joanne Lachica, Jaijeet Singh, Ali Ghareeb, James Bayne, Melissa Grant-Peters, Sonia Garcia-Ruiz, Zhongbo Chen, Samuel Rodriques, Dilan Athauda, Emil Gustavsson, Sarah A. Gagliano Taliun, Christina Toomey, Regina H. Reynolds, George Young, Stephanie Strohbuecker, Tom Warner, Dmitri A. Rusakov, Rickie Patani, Clare Bryant, David A. Klenerman, Sonia Gandhi, Mina Ryten

**Author notes:** Joint Authors (Karishma D’Sa, Minee L. Choi, Aaron Z. Wagen, Sonia Gandhi, Mina Ryten).

## Abstract

RNA editing is a post transcriptional mechanism that targets changes in RNA transcripts to modulate innate immune responses. We report the role of astrocyte specific, ADAR1 mediated RNA editing in neuroinflammation in Parkinson’s disease. We generated hiPSC-derived astrocytes, neurons and co-cultures and exposed them to small soluble alpha-synuclein aggregates. Oligomeric alpha-synuclein triggered an inflammatory glial state associated with TLR activation, viral responses, and cytokine secretion. This reactive state resulted in loss of neurosupportive functions, and the induction of neuronal toxicity. Notably, interferon response pathways were activated leading to upregulation, and isoform switching of the RNA deaminase enzyme, ADAR1. ADAR1 mediates A-to-I RNA editing, and increases in RNA editing were observed in inflammatory pathways in cells, as well as in post-mortem human PD brain. Aberrant, or dysregulated, ADAR1 responses and RNA editing may lead to sustained inflammatory reactive states in astrocytes triggered by alpha-synuclein aggregation, and this may drive the neuroinflammatory cascade in Parkinson’s.

## Introduction

Parkinson’s disease (PD) is a progressive neurodegenerative condition characterised by the accumulation of intraneuronal aggregates of alpha-synuclein (α-syn) through the brain (*1*). Astrogliosis has been reported in post-mortem Parkinson’s brain (*2–7*), and in rodent models of synucleinopathy (*8*, *9*), and astrocytes accumulate α-syn inclusions (*10*), raising a role for astrocytes, and astrocyte-mediated immune cascades (*11*), in triggering or driving PD pathogenesis. Astrocytes are an abundant glial cell, essential for neuronal function, and survival through the maintenance of CNS homeostasis via modulation of neurotransmitters and synapse formation, ionic balance, and maintenance of the blood-brain barrier (*12*). Reactive astrogliosis defines a process whereby, in response to pathology, astrocytes undergo changes in transcriptional regulation, biochemical, morphological, metabolic, and physiological remodeling, which ultimately result in a switch from resting to reactive states. The nature of the underlying stimulus (neuroinflammatory vs ischaemic) has defined certain reactive astrocyte states into neurotoxic (so called A1, which promote lipid mediated neuronal death via activated microglia) and neuroprotective (so called A2, which promote neuronal survival and regeneration through neurotrophic factors) (*13*). Reactive transformation can be associated with loss of neurosupportive and homeostatic functions, reduced synapse formation (*14*), alterations in glutamate uptake and recycling (*15*), and dysregulated calcium signalling (*16*). Simultaneously reactive states are associated with the gain of neurotoxic properties (*17*).

Focusing on PD, there is evidence that A1 astrocytes have been shown to infiltrate brain regions associated with PD, including the striatum(*13*). Astrocytes in these regions express the highest regional levels of immune mediators, such as toll-like receptor 4 (TLR4) and myeloid differentiation primary response 88 (MyD88), correlating with PD pathology (*18*). Significantly, inhibiting the formation of these A1 astrocytes by blocking microglia-astrocytic cross talk is neuroprotective in mouse models of PD (*19*). Reactive astrocyte substates are likely to be diverse beyond these two states (*14*, *20*), and involve several exogenous and endogenous triggers such as damage-associated molecular patterns (DAMPs) or proteostatic stresses induced by misfolded proteins. Compared to control cell lines, human astrocytic models of PD release increased amounts of pro-inflammatory cytokines in response to α-syn (*21*). Moreover, in response to α-syn fibrils, iPSC-derived astrocytes can assume an antigen presenting function, with upregulation of major histocompatibility complex (MHC) genes and relocation of HLA molecules to the cell surface to present α-syn fibril peptides to neighbouring cells (*21*). This finding, replicated in post-mortem brain samples (*22*), supports the idea that astrocytes may mediate the immune response in the brain.

We have recently reported that physiological concentrations of oligomeric α-syn, trigger a TLR4-dependent inflammatory response in murine primary astrocytes (*18*). The small soluble hydrophobic and beta sheet rich aggregates, or oligomers of α-syn are known to be neurotoxic, and trigger cell selective processes that are specific to the structural conformation of the protein (*23–26*).

Sub-state modelling has been achieved in human iPSC-derived astrocytes and co-cultures (*27*), which enable capture at a molecular level of cell autonomous and non-cell autonomous cascades in pathology. We investigate how human iPSC-derived astrocytes respond to α-syn oligomers (αsyn-O), which downstream response pathways are activated, and how this reactivity affects neurons. We integrate bulk and single-cell transcriptomic, functional and biophysical approaches in five lines of human iPSC-derived (hiPSC-derived) astrocytes (3 in-house and 2 commercial lines), both alone and in co-culture with neurons, to define the molecular response of astrocytes to misfolded α-syn. Finally, identifying innate immune pathways of interest, we explore those pathways in post-mortem brain samples from those with PD.

## Results

### Generation and functional characterisation of hiPSC-derived astrocytes and neurons alone or in co-culture

We generated hiPSC-derived astrocytes from 5 healthy donors, and cortical neurons from a sixth healthy donor (**Supp table 1**) (*28*). We compared the cellular response to αsyn-O in astrocyte only, neuron only and astro-neuronal cultures (**Figure 1a**). Highly enriched cultures of cortical astrocytes in feeder-free conditions were generated from three in-house lines using an optimised small molecule serum free protocol, and two were purchased commercially, derived through a proprietary serum free protocol (**Figure 1b**). We used the Shi et al., 2012 protocol (**Figure 1b**) to generate highly enriched (>90%) and functional hiPSC-derived cortical neurons from neural precursor cells (*24*, *25*).

**Fig 1.**
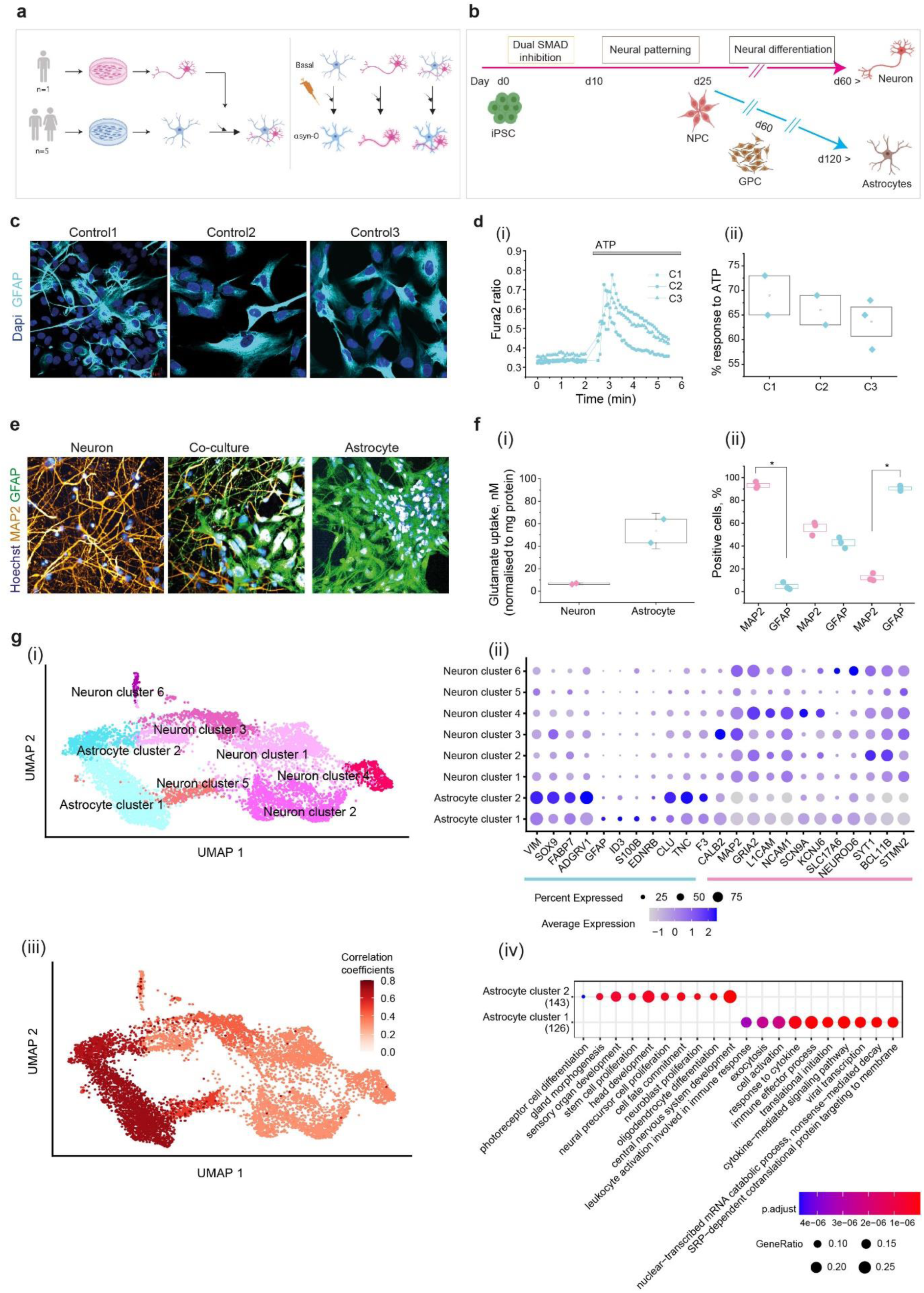
Schematic illustrations of experimental paradigm and the differentiation protocols. (a) hiPSC-derived from five healthy individuals are differentiated into astrocytes (in-house lines: C1, C2, C3, commercial lines from ACS: C4, C5) and from one individual into neurons. (b) Differentiation of cortical region-specific astrocytes was performed, modifying established protocols.(*76*, *77*) Neural precursor cells (NPCs) using an established protocol (Shi et al., 2012) were differentiated into either neurons and GPC (glial precursor cells) after 30 days from NPC, then further differentiated into mature astrocytes. (c) Immunocytochemistry images show that three lines of hiPSC-derived astrocytes express the astrocytic marker, GFAP. (di-ii) There is a calcium response to APT in iPSC-derived astrocytes. Representative traces of calcium (di) and the percentage of cells (dii) in response to ATP. (e) hiPSC-derived astrocyte enables uptake of glutamate (Glutamate Assay Kit, (ab83389/K629-100, Abcam). (fi-ii) Composition of hiPSC-derived neuron and astrocyte co-culture assessed using MAP2 (neuronal marker) and GFAP (astrocytic marker) immunocytochemistry together with representative images of a neuronal, astrocyte and co-culture (fi) and the quantification (fii). (g) (i)UMAP plot showing the clustering of the integrated dataset using the cells from all the samples (basal and αsyn-O treated astrocytes, neurons, co-culture samples). (ii) Dot plots showing the expression of the astrocyte and neuron marker genes in the clusters identified in the single-cell RNA-Seq data. (iii) UMAP overlaid with the correlation coefficients, showing the correlation of the 2 astrocyte clusters with the astrocytes from Leng et al.(*27*) (iv) GO terms associated with the genes up-regulated in Astrocyte clusters 1 and 2.

Immunolabelling with GFAP, an astrocyte-specific marker demonstrated that cultures contained >90% mature astrocytes (**Figure 1c**). hiPSC-derived astrocytes also displayed cytosolic calcium responses on application of ATP as assessed using Fura-2 (**Figure 1d**) (*29*). All lines demonstrated appropriate Na+ dependent uptake of glutamate from the extracellular space, one of the major astrocytic functions that indicates functional activity of Excitatory Amino Acid Transporter 1/2 in hiPSC-derived astrocytes (**Figure 1f(i)**). Together, the ATP-dependent calcium responses and glutamate uptake confirmed generation of functionally mature astrocytes.

To investigate how astrocytes affect neuronal function, we generated co-cultures by plating hiPSC-derived cortical astrocytes and neurons in a 1:1 ratio, and cultured for at least 3 days prior to use. The composition of co-cultures was assessed by immunocytochemistry using both MAP2, a neuronal marker, and GFAP, an astrocytic marker (**Figure 1e, 1f(ii)**). The co-cultures contained 55.9 ± 3.4% MAP2 positive cells and 42.9 ± 2.7% GFAP positive cells, while neuronal cultures contained 93.2 ± 1.7% MAP2 positive cells and astrocytic cultures contained 90.5 ± 1.3% GFAP positive cells.

In order to generate oligomers of α-syn, human recombinant monomeric WT α-Syn was aggregated in the dark at 37°C and 200 r.p.m., for ∼7–8 hours. At this time point, the mixture consists of 99% monomeric α-syn, and 1% oligomeric α-syn, with oligomers being small, soluble, and rich in beta sheet structure as characterised previously (*24*, *25*, *30*).

Whole cell patch-clamp recording of neurons was performed in neuron only and astro-neuronal cultures to assess the electrophysiological properties and excitability of the cells. In neuron only cultures, neurons (∼100 DIV) displayed a relatively depolarised resting membrane potential (*V*rest) compared with age-matched cells in co-cultures (–45.9 ± 2.8 mV, n = 33 vs. –56.0 ± 2.1 mV, n = 34, p = 0.0053, respectively, **Supp figure 1(a,b)**). No significant difference was observed in neuronal capacitance across culture types (*C*_m_: 44.8 ± 3.5 pF, *n* = 35 in cultures and 43.7 ± 3.8 pF, *n* = 40 in co-cultures, *p* = 0.843; (**Supp figure 1c**). Input resistance (**Supp figure 1d**) and the time constant (**Supp figure 1e**) were significantly altered in astro-neuronal cultures. In neuron only cultures, neurons generated a single action potential (AP) in response to current injection, whereas neurons in astro-neuronal cultures generated a train of induced APs in either step-wise depolarising protocol or slow-injecting ramp current (**Supp figure 1f**). The parameters of AP spike (threshold, spike amplitude, kinetics) also confirmed that co-cultures altered the neuronal performance. Neurons were more excitable in co-cultures, as a lesser current was required to bring neurons to drive firing ( **Supp figure 1g-j**).

### Single-cell RNA-sequencing of astrocyte only, neuron only and astro-neuronal cultures

We used RNA-sequencing to further characterise hiPSC-derived astrocytes and neurons. After 120 days of differentiation all cells were harvested with and without αSyn-O stimulation, each with a technical replicate resulting in 44 samples. Using the ‘in house’ protocol, cultures were sequenced with single-cell technology (astrocyte only, neuron only and astro-neuronal cultures, with and without αsyn-O). Across the integrated single-cell dataset we identified 8 cell clusters, which include both astrocytic and neuronal subtypes. Cell types were assigned using a set of previously published and curated marker genes (*31*, *32*). Based on the expression of these marker genes, we identified two astrocytic and six neuronal clusters (**Figure 1g (i,ii)**). Given that hiPSC-derived astrocytic profiles have been previously well characterised by Leng and colleagues (*27*) we initially assessed our clusters for correlations in gene expression globally with the reported iAstrocytes transcriptomic profiles and found they were highly correlated with both of our astrocyte clusters (r = 0.71 and 0.64 ) (**Figure 1g(iii)**), subsequently referred to as astrocyte cluster 1 (AC1) and 2 (AC2).

Given the growing literature on astrocytic subtypes, we investigated the clusters further. Focusing specifically on genes differentially expressed between the clusters (AC1 and AC2), we identified 300 genes of interest (at FDR< 5% and >2-fold change in expression) of which 129 were more highly expressed in AC1 and 171 genes which were more highly expressed in AC2 (**Supp table 2**). Interestingly, gene set enrichment analysis of these differentially expressed genes, identified immune and cytokine-related terms amongst genes more highly expressed in AC1. Conversely, genes more highly expressed in AC2 showed enrichment for terms related to morphogenesis, regulation of development and differentiation (**Figure 1g(iv), Supp table 3**). Furthermore, we noted that the pattern of gene expression in the AC2 cluster appeared to resemble that described for neuroprotective astrocytes (**Supp figure 2a**). Thus, we identified 2 subtypes of astrocytes, AC1 and AC2, which differed in terms of their inflammatory and protective transcriptomic profiles.

### hiPSC-derived astrocytes are reactive to αsyn-O and secrete cytokines

Next, we examined whether hiPSC-derived astrocytes take up αsyn-O using a Fluorescence Resonance Energy Transfer (FRET) biosensor which enables visualisation of oligomers in cells (*25*) We treated hiPSC-derived astrocytes with two populations of fluorescently tagged α-syn (AF488-& AF594-tagged A53T α-syn monomers, total 500 nM) for 5 days, and then measured the intracellular accumulation of α-syn (‘total α-synuclein’) based on the intensity of AF594 through direct excitation with 594 nm irradiation. The formation of oligomer (‘FRET’) was visualised via the presence of signal from the acceptor fluorophore (AF594) after excitation of the donor fluorophore (488 nm irradiation) (**Figure 2a**). Using this approach, we found total α-syn uptake was higher in hiPSC-derived astrocytes than neurons, while similar levels of de novo aggregates (FRET) were detected in both cell preparations (**Figure 2b**).

**Fig 2.**
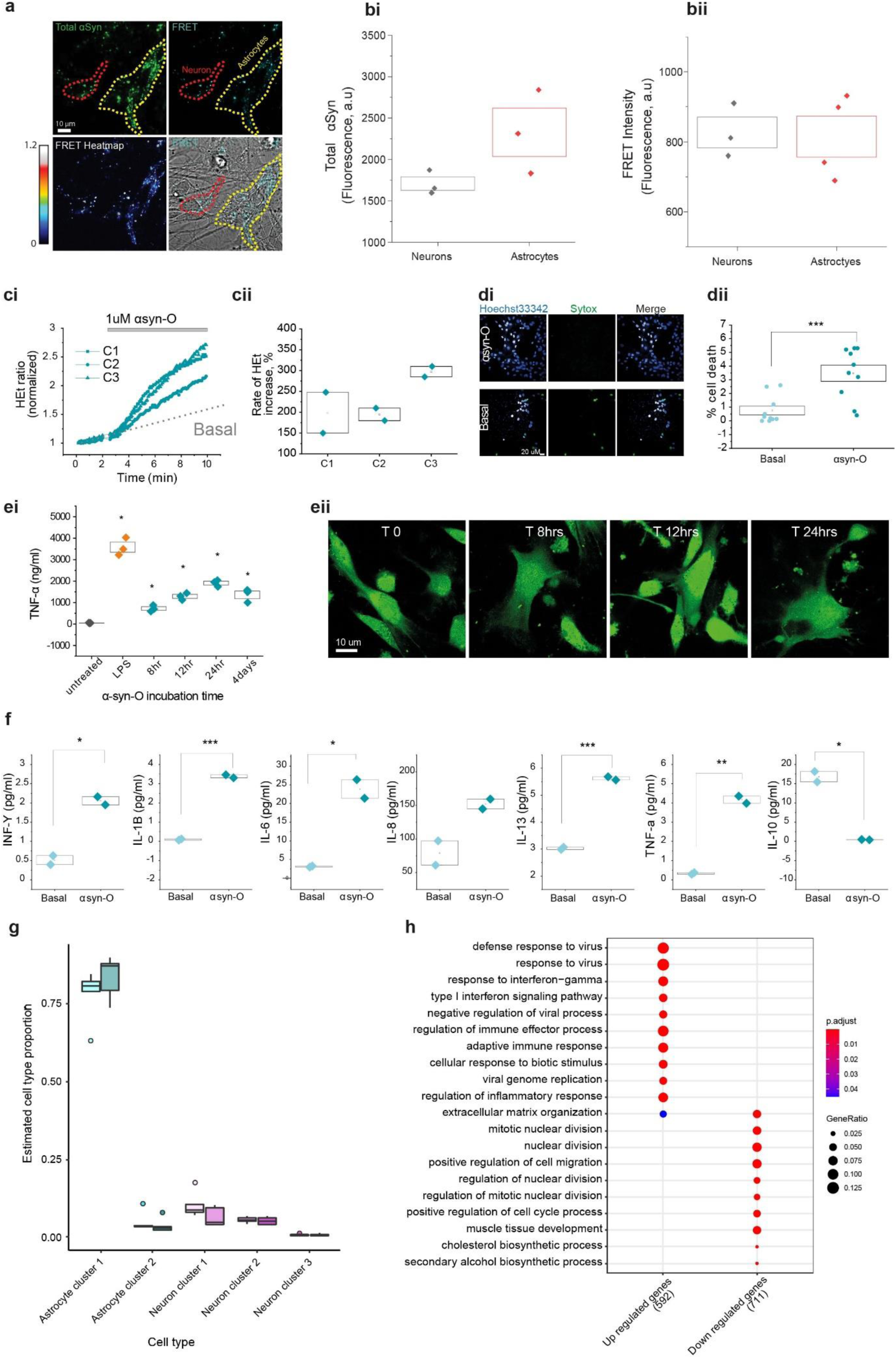
Oligomer treatment of astrocytes induces an inflammatory state. (a) Uptake of monomeric species of labelled α-syn (A53T monomer) in astrocytes (with yellow dotted lines) and neurons (with red dotted lines) was detected, and the formation of oligomers inside cells confirmed using the FRET biosensor. (bi-ii) There is no difference in the formation of oligomeric species despite the higher uptake of total α-syn in astrocytes compared to neurons (n = 3 independent inductions). (ci-ii) Application of αsyn-O induces the overproduction of reactive oxygen species (ROS). (di-ii) Cell death induced by αsyn-O is detected in astrocytes at low levels. (ei) αsyn-O treated astrocytes release cytokine responding to αsyn-O (measured by variable incubation time and the representative images). Cytokine release is time-dependent, with the level of TNF-α highest after 24hr incubation with αsyn-O. (eii) Morphological changes were traced using Fluo4 across the same time course as the cytokine release measurements. (fi-vii) αsyn-O treated astrocytes induce an inflammatory state by releasing a range of cytokines responding to αsyn-O (basal IL-13: 3.02 ± 0.04 pg/ml, αsyn-O treated 5.62 ± 0.05 pg/ml, IL-6: basal 3.11 ± 0.13 pg/ml, αsyn-O treated 23.8 ± 2.4 pg/ml, IL-8: basal 79.11 ±18.07 pg/ml, αsyn-O treated: 152.01 ± 7.29 pg/ml, IL-1B: basal 0.955 ± 0.03011, αsyn-O treated 3.38 ± 0.0772, INF-Y: basal 0.51 ± 0.11, αsyn-O treated 2.05 ± 0.1 pg/ml, TNF-α: basal 0.33 ± 0.0414, αsyn-O treated 4.16 ± 0.187) (g) Cell type proportions derived from deconvolution using Scaden. The plot shows the cell type proportion for each of the cell types in the bulk astrocyte samples, basally and with αsyn-O treatment, as predicted by Scaden. Cell types with proportions < 1% were excluded. (h) The top 10 GO terms associated with up and down-regulated differentially expressed genes at FDR < 5% and with at least >= 2-fold change in expression in the astrocytes basally vs with αsyn-O treatment.

αsyn-O exposure can cause cell toxicity, and excessive ROS generation in primary astrocytes (*18*). To determine whether hiPSC-derived astrocytes show similar responses, we assessed ROS production using dihydroethidium (DHE) dye which allowed us to robustly measure the rate of oxidation of the dye by cellular superoxide production (*33*). We found that ROS production significantly increased compared to basal levels (normalised to 100%, **Figure 2c**) in hiPSC-derived astrocytes after the application of αsyn-O.

Since our previous study showed that αsyn-O treatment triggers an inflammatory response in primary astrocytes with associated increases in cytokine release (*18*), we assessed this phenomenon in hiPSC-derived astrocytes. Following treatment of hiPSC-derived astrocytes with a range of αsyn-O concentrations (100nM – 2uM) overnight, we collected the media and assessed the cytokine profiles using an MSD (V-PLEX Proinflammatory Panel 1) electrochemiluminescence assay kit. We found that αsyn-O treatment of hiPSC-derived astrocytes consistently induced a significant increase in the secretion of a variety of cytokines compared to untreated cultures. Thus, we demonstrated that αsyn-O exposure induces hiPSC-derived astrocytes to become pro-inflammatory (**Figure 2e,f**).

### Oligomer treatment of hiPSC-derived astrocytes triggers anti-viral inflammatory responses

Next, we used bulk RNA-sequencing to investigate hiPSC-derived astrocyte responses to αsyn-O treatment in more detail. Given that we identified two major astrocytic clusters (AC1 and AC2), we determined whether αsyn-O changes the relative proportions of these cell clusters. With this in mind, we used the tool Scaden (*34*) together with our scRNA-seq data (Materials and methods) to estimate cluster proportions in bulk RNA-Seq data across the whole dataset, noting a high correlation in cell type proportion estimates based on single-cell and Scaden-derived data (**Supp figure 2b**). This approach was based on the colocalization of all astrocytic samples within exploratory principal component analyses based on bulk RNA-seq data, so suggesting that the astrocyte clusters we identified in a subset of samples were in fact present in all. While we found no significant change in astrocyte subtype proportion between basal and αsyn-O treated astrocyte cultures (**Figure 2g, Supp table 4**), significant changes in gene expression and splicing were observed.

We identified 2004 genes which were significantly differentially expressed (8.17% at FDR < 5% and at least 2-fold change in expression) following treatment of hiPSC-derived astrocytes with αsyn-O, of which 917 were up-regulated and 1087 were down-regulated in the treated astrocytes (**Supp table 5**). Importantly, these up-regulated genes with at least 2-fold change in expression were enriched for those implicated in viral responses, including “defence response to virus”, “response to interferon-gamma”, “type I interferon signalling pathway” (**Figure 2h, Supp table 6**). These results were highly consistent with the functional data demonstrating a robust cytokine response to αsyn-O treatment. Since splicing analyses have been shown to provide distinct biological information (*35–37*), this form of analysis was used to further characterise hiPSC-derived astrocyte responses to αsyn-O. We identified 707 significant differentially spliced intron clusters corresponding to 590 genes (FDR < 0.05, |ΔPSI|≥ 0.1) with significant enrichment for cytoskeletal terms, potentially reflecting the observed morphological changes in astrocytes with oligomer treatment (**Supp figure 3 & supp tables 7,8,9**). Morphological changes induced by αsyn-O are evaluated through astrocyte segmentation, followed by quantification of GFAP pixel area and intensity based on the distance from the nuclear membrane. This approach assesses both morphological polarity and intensity gradients (*38*, *39*), providing a detailed understanding of the spatial distribution and intensity changes in astrocytic morphology thus enabling the tracking of reactive astrocytic morphology upon αsyn-O stimulation.

### hiPSC-derived astrocytes maintain inflammatory states on exposure to αsyn-O in co-culture

First, we studied the functional effects of αsyn-O treatment on astro-neuronal cultures. Media collected from the samples were used to measure a range of cytokines secreted from the cells using an MSD (V-PLEX Proinflammatory Panel 1) electrochemiluminescence assay kit. αsyn-O induced more secretion of a variety of cytokines compared to untreated co-cultures demonstrating inflammatory activation of the co-culture (**Figure 3a**).

**Fig 3.**
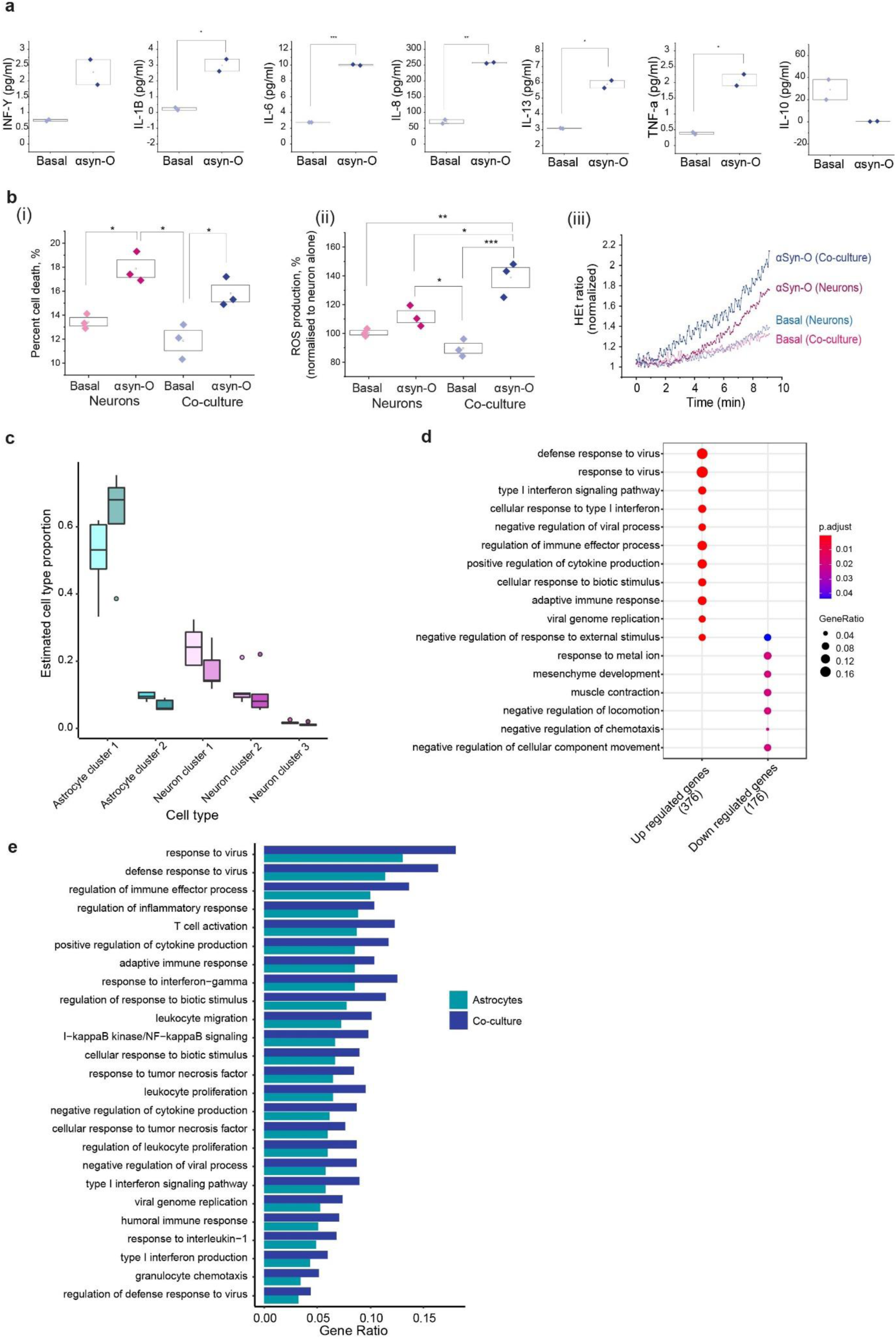
Co-culture setting provides evidence of the astrocytes becoming more inflammatory. (a) Effect of αsyn-O on cytokine release in the co-cultures (basal IL-13: 3.091 ± 0.0139 pg/ml, αsyn-O treated 5.88 ± 0.238 pg/ml, IL-6: basal 2.75 ± 0.00174 pg/ml, αsyn-O treated 10.026 ± 0.0593 pg/ml, IL-8: basal 71.09 ± 6.107 pg/ml, αsyn-O treated 257.2 ± 1.58 pg/ml, IL-1B: basal: 0.231 ± 0.0835 pg/ml, αsyn-O treated 2.99 ± 0.381 pg/ml, INF-Y: basal 0.392 ± 0.0329, αsyn-O treated 2.078 ± 0.180 pg/ml, TNF-α: basal 0.392 ± 0.0329, αsyn-O treated 2.078 ± 0.1801) (bi-iii) αsyn-O induces toxicity in neurons alone and in co-culture with astrocytes, and induces increased levels of ROS in neurons alone and in co-culture. (c) Plot showing the cell type proportions estimated by Scaden from the single cell data in the co-culture samples, basally (lighter shade) and with αsyn-O treatment (darker shade). There is a significant difference between the cell type proportions of astrocyte cluster 1 and 2 in the co-culture with αsyn-O treated compared to basal. (d) GO terms associated with the up- and down-regulated differentially expressed genes with at least >= 2-fold change in expression in the co-culture αsyn-O treated vs basal (e) Visualising gene ratios of the enrichments observed in the up-regulated differentially expressed genes of the astrocytes (cyan) compared to that of the co-culture (blue).

Previously we have demonstrated that αsyn-O induced an increased level of neuronal death and oxidative stress (*18*, *24*, *25*). Using live-cell imaging, we showed that activated astro-neuronal cultures were associated with higher levels of cell death (**Figure 3b(i)**), and higher levels of ROS production (**Figure 3b(ii, iii)**). Furthermore, patch-clamp recordings performed in co-cultures treated with αsyn-O demonstrated a drop in the *V*rest after treatment (**Supp figure 4b**) and increased input resistance (**Supp figure 4c**) in neurons. Neurons also had impaired firing, with a dramatically changed AP waveform. The threshold for AP spike generation was depolarised (**Supp figure 4d,e**), the amplitude was reduced (**Supp figure 4f**), and the spike was significantly extended compared with control astro-neuronal cultures. Thus the proteinopathy appeared to induce an activated inflammatory state of astrocytes, that is associated with loss of the previous neuronal supportive function seen with resting astrocytes. Additionally there are toxic gain-of-function effects in both astrocytes and neurons in co-culture, such as induction of oxidative stress, altered excitability, and neuronal cell death.

As before, we also combined single-cell and bulk RNA-sequencing analyses to assess the impact of αsyn-O treatment on cell subtype proportions, gene expression and splicing. We observed a significant increase in the proportion of AC1 (inflammatory) relative to AC2 (neuro-protective) astrocytes in αsyn-O treated as compared to co-cultures basally (**Figure 3c, Supp table 4**). We noted that there was no significant difference observed in the cell type proportions of AC1 and AC2 in astrocyte only cultures on αsyn-O treatment (**Supp table 4**). Differential gene expression analysis following correction for changes in predicted cell type proportions, identified 774 genes (3.46%, FDR < 5% and at least 2-fold change in expression) with 509 up-regulated and 265 down-regulated in the αsyn-O treated as compared to the co-cultures basally (**Supp table 5**). Similar to the findings on αsyn-O treated astrocytes only, the up-regulated genes with at least 2-fold change in expression were highly enriched for immune response terms (**Figure 3d, Supp table 10**). As observed in the astrocyte only cultures, the terms highlighted were associated with viral infections, such as “defence response to virus” and “type 1 interferon signalling pathway”. Furthermore, we noted that gene enrichment ratios for immune-related GO terms were consistently higher in αsyn-O treated co-cultures compared to astrocyte only cultures, suggesting a more prominent inflammatory response to oligomer treatment in co-cultures (**Figure 3e, Supp table 11**).

Again, analysis of differential splicing identified distinct biological processes that were related to structural organisation of cells and physical cell interactions, in contrast to gene level expression signals. We identified 502 differentially spliced intron clusters in 414 genes (FDR < 0.05, |ΔPSI|≥ 0.1), with the genes enriched for terms relating to junction assembly and synapse (**Supp table 7**). Finally, we assessed differentially expressed and differentially spliced genes for evidence of enrichment for genes genetically associated with either Mendelian forms of early onset PD and Parkinsonism, or complex PD (*40*, *41*). While we did not see any enrichment of Mendelian genes in astrocyte only cultures, there was significant enrichment amongst all genes that were differentially expressed or differentially spliced (**Supp table 12**) in astro-neuronal cultures. Overall, this suggests that this model is related to PD pathogenesis and uncovers pathways related to PD causation.

### Oligomer treatment triggers *ADAR* expression and a change in isoform use

αsyn-O treatment of both astrocyte only and astro-neuronal cultures resulted in the activation of pathways most commonly associated with responses to viruses. In both cases, we noted significant increases in the expression of genes such as *MDA5*, *RIG1* and *TLR3* that can sense viral RNA (double-stranded RNA or Z-RNA)(*42–44*) and activate the release of IFN and cytokines (**Figure 4a, Supp figure 5b**). This in turn is known to trigger the up-regulation of a range of genes, including *OAS1*, *PKR* and *ZBP1* to degrade viral RNA, inhibit translation and drive necroptosis respectively, and indeed this up-regulation was identified in αsyn-O treated cultures (**Figure 4a, Supp figure 5b**).

**Fig 4.**
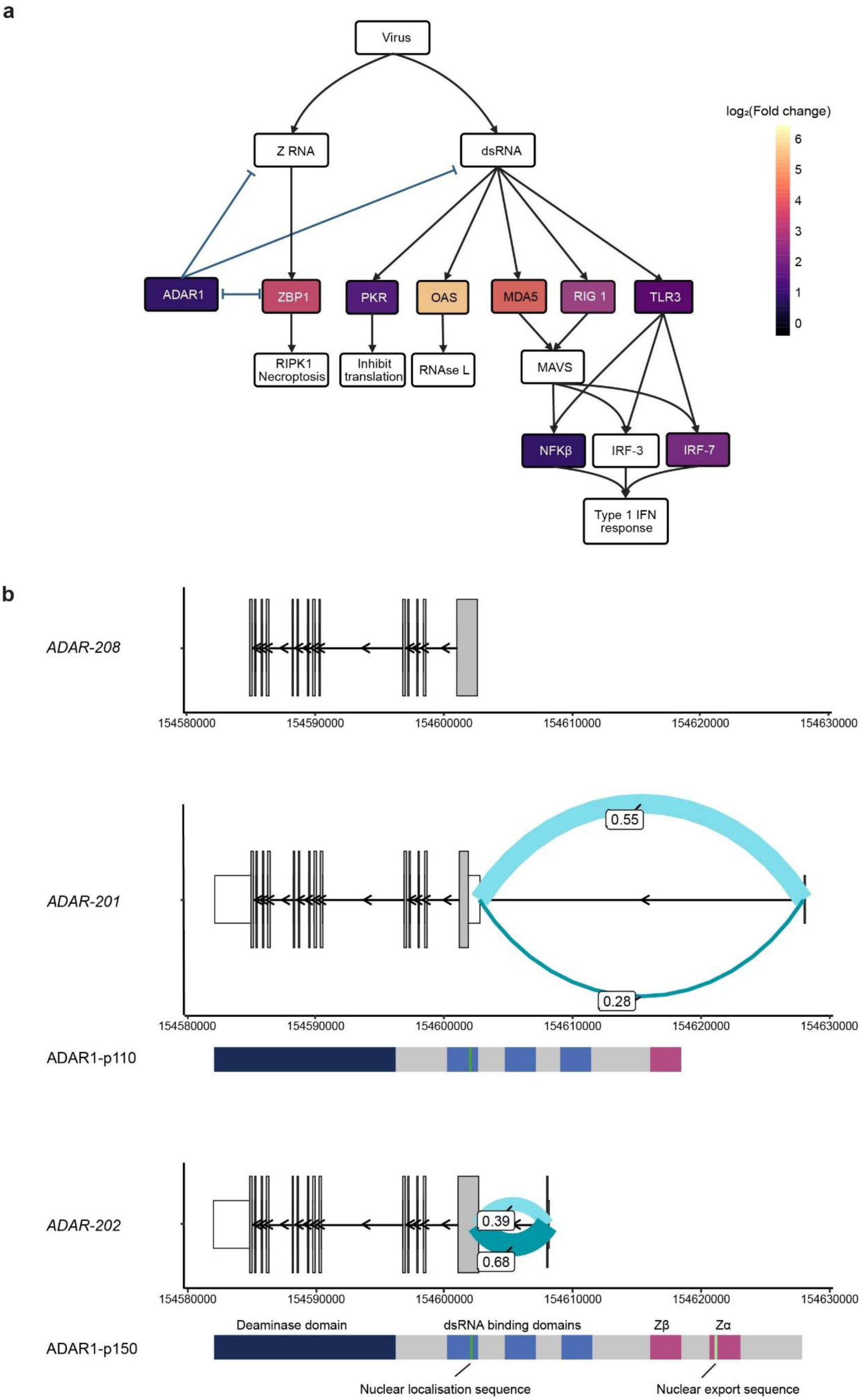
Type 1 IFN response leads to activation of *ADAR*. (a) Model summarizing the role of *ADAR* in regulating the innate immune response to dsRNA with the log_2_(Fold change) of the genes that are significantly differentially expressed (at FDR < 5%) in the astrocytes on αsyn-O treatment. (b) Plot showing the significantly differentially spliced junctions in *ADAR* in the astrocytes on αsyn-O treatment, with the transcripts and protein isoforms. (Co-culture: **Supp figure 5a, b**).

However, these processes also undergo negative regulation by ADAR, an enzyme which deaminases adenosines on double stranded RNA to inosines (*45*). This conversion reduces the activation of dsRNA sensors by disrupting RNA self-complementarity and so favouring the formation of single-stranded RNA forms, as well as through other direct and indirect modulation of pro-inflammatory pathways (*46–49*). In fact, it is known that biallelic pathogenic variants in ADAR that reduce its editing activity, result in excessive release of interferons and tissue damage, and present as Aicardi Goutieres syndrome (*50*). With this in mind, we noted that αsyn-O treatment of both astrocyte and astro-neuronal cultures resulted in significant increases in *ADAR* expression (**Figure 4a, Supp figure 5c**) of 1.86 and 1.82 fold in the cultures respectively.

Furthermore, αsyn-O treatment resulted in significant differences in transcript use (**Figure 4b, Supp figure 5a**) as detected through splicing analyses. More specifically, we noted a decrease in the use of an exon-exon junction specific to *ADAR*-201 (1:154602627-154627854:-) encoding the ADAR p110 isoform, and a relative increase in the usage of an exon-exon junction (1:154602627-154607991:-) specific to *ADAR*-202 encoding the p150 isoform, which is already known to be under the control of an IFN-sensitive promoter (**Figure 4b, Supp figure 5a**). Thus, the functional and transcriptomic analyses of astrocyte only and astro-neuronal cultures, suggest that αsyn-O treatment triggers the increased expression of *ADAR* and an increase in the use of the cytoplasmic p150 isoform.

### αsyn-O treatment increases A-to-I editing in astrocyte-containing samples

We postulated that changes in *ADAR* expression and its isoform use would result in both an increase in A-to-I editing rate and a change in the distribution of A-to-I editing sites. The latter, would be expected as a consequence of the different properties of ADAR’s two major isoforms, with the p110 exclusively found in the nucleus and the p150 being largely cytoplasmic (*51*). To investigate this we used the high-depth bulk RNA-seq data we generated across all cultures to identify editing sites, in each case comparing to the reference genome.

Focusing on basal conditions, we detected 78,547 editing sites in astrocyte only cultures, 103,340 sites in the neuron only cultures, and 100,740 sites in the astro-neuronal cultures (**Figure 5a**). Consistent with the known higher levels of editing in neurons (*52*), we found that the median baseline editing rate at a given site was higher in neuron only (0.208) than in astrocyte-containing cultures (0.054 in astrocyte only, 0.056 in astro-neuronal cultures). Given that astro-neuronal cultures were composed of ∼50% neurons, our findings suggest that in the presence of astrocytes editing rates in neurons are lower.

**Fig 5.**
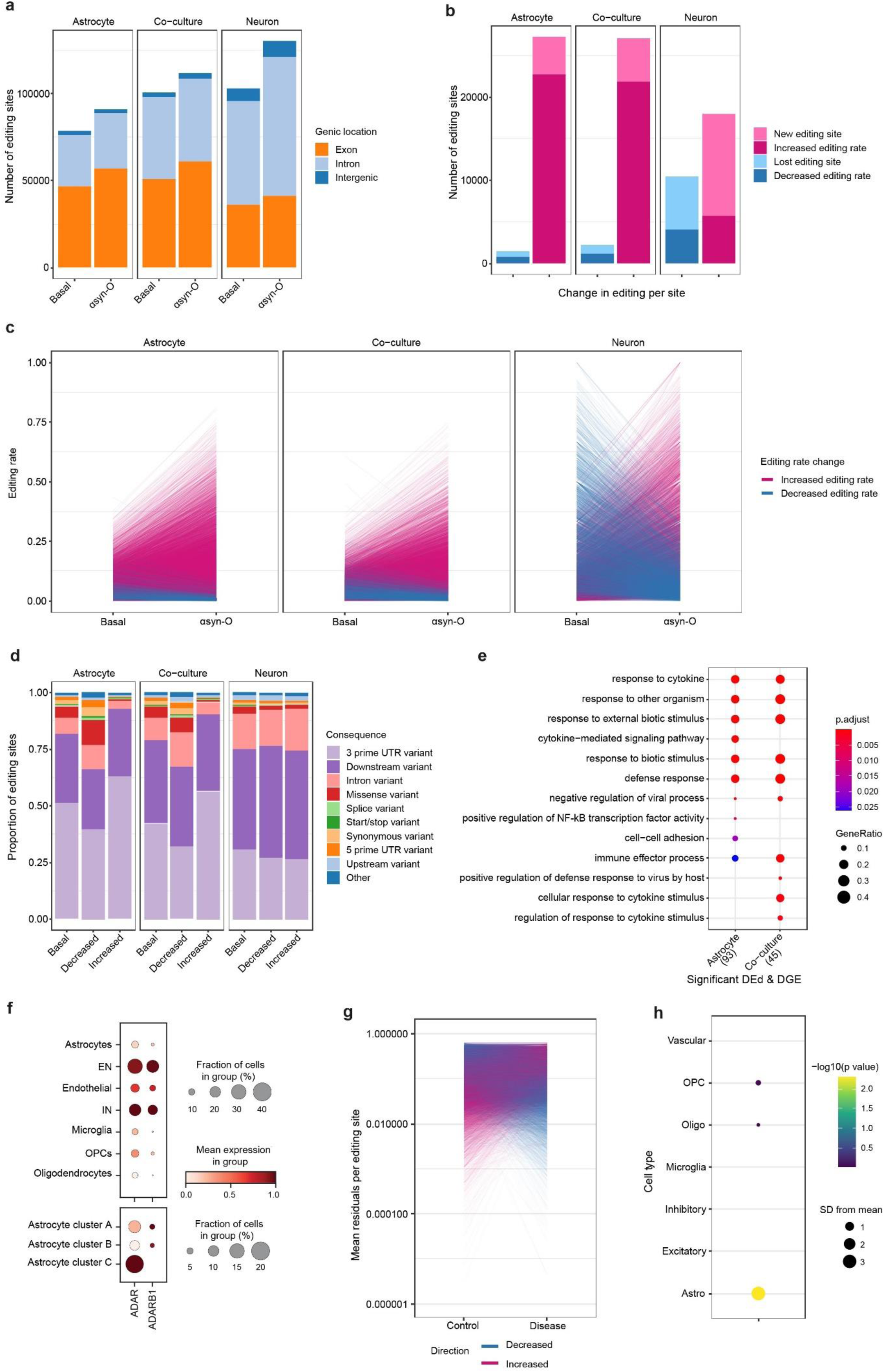
*ADAR* induced in A to I editing in astrocytes and co-cultures with αsyn-O treatment. (a) Number of editing sites in each sample, showing genomic locations (b) Number of differentially edited sites when comparing αsyn-O treated to samples basally in each culture. X-axis shows the decreased editing (blue, including lost) and increased editing (pink, including new). (c) Change in editing rate in basal vs αsyn-O treated samples in each culture. (d) Consequences of editing sites at baseline, and in differentially decreased or increased sites across cultures. (e) GO terms associated with the significantly differentially edited and differentially expressed (FDR < 5% and FC >=2) in the astrocytes and co-culture αsyn-O, treated vs basal. (f) Expression of *ADAR* and *ADARB1* in single nuclear post-mortem brain RNA. The top panel shows expression by cell-type, and the bottom shows expression within astrocytic clusters. Astrocyte cluster A is defined by expression of *ADGRV1* and *SLC1A2*; Astrocyte cluster B by *GFAP*, *S100B*, *AQP4* and Astrocyte cluster C by *VIM*, *SOX9* and *FOS*. (g) Mean residuals per editing site, after correction for covariates, in controls and diseased PD post-mortem brain samples. (h) Expression weighted celltype enrichment analysis of genes with 50 greatest increase in mean editing rate. Abbreviations: EN - excitatory neurons. IN - inhibitory neurons. OPC - oligodendrocyte progenitor cells. Supplementary Figures-Supp fig 1 to Supp fig 7

This analysis also revealed differences in the distribution of editing sites across cell cultures. In astrocyte only cultures, 50.3-59.5% of editing sites were in exonic regions, with the majority of these located in the 3’UTR (49.6% and 40.9% of editing sites in astrocyte only and astro-neuronal cultures, respectively **Figure 5d**). By contrast, in the neuron only cultures we found that just 25.4% of sites were located in the 3’UTR. Finally, consistent with the known molecular function of *ADAR* we found that irrespective of genic location or culture type, the majority of editing sites (83.6 – 86.6%) were located within repeat regions, of which the vast majority were in Alu regions (93.2 – 94.5%) (**Supp Figure 6c**).

As predicted, αsyn-O treatment generated an increase in the number of editing sites in all cultures (15.8% in astrocyte only, 26.2% in neuron only, and 11.1% in astro-neuronal cultures), which was highly significant (chi-squared p-value of <2×10^-16^ for increased exonic proportion in all cases). Similarly, measurement of editing rates at each site demonstrated both a marked increase in the number of new editing sites and editing rate in astrocyte and astro-neuronal cultures, but not neuron only cultures (**Figure 5b**). In contrast, across all cultures relatively few editing sites were lost or had a decrease in editing rate.

Furthermore, consistent with a change in transcript usage, αsyn-O treatment was associated with a change in the distribution of editing sites. In astrocyte-containing samples, the sites with increased editing rates were significantly more likely to be in 3’UTRs than sites with decreased editing (p-value < 2×10^-16^ in both astrocyte only and astro-neuronal cultures), while in neuronal monocultures the proportion of sites in 3’UTRs did not change significantly (p-value = 0.17). A similar pattern was observed when exploring the biotype (as defined by Ensembl VEP 93.5) of the transcripts containing a given editing site, with sites identified to have an increase in editing rate in astrocyte-containing cultures being more likely to be located within protein coding transcripts (p-values < 8.2×10^-8^ and 2×10^-16^ in astrocyte only and astro-neuronal cultures respectively, **Supp figure 6b**). Taken together, these results show that αsyn-O treatment is associated with an increase in the number of editing sites and differential editing rate in astrocyte-containing cultures.

Noting that RNA editing can influence gene expression through effects on mRNA stability, we explored the relationship between transcript editing, gene expression and gene function (*53*). We began by identifying all genes that both contained sites with significant differential editing on exposure of cells to αsyn-O treatment, and which had significant differences in gene expression in the same conditions. We found that there was a significant overlap in genes that were differentially edited and differentially expressed in the astrocyte only and astro-neuronal cultures following αsyn-O treatment (Fisher’s exact test p-value for astrocytes 8.54E-4, co-culture 2.39E-2). Focusing on this gene set, namely genes that were both differentially edited and expressed, we found a significant enrichment for the terms linked to viral infection and immune response in both the astrocyte-containing cultures (**Figure 5e**, **Supp table 13**).

### A-to-I RNA editing is increased in post-mortem PD brains

While A-to-I editing in the human brain is well-recognised and perhaps best characterised in neurons, the molecular machinery for editing is present in all the major cell types. In humans A-to-I editing is catalysed not only by ADAR1 (*ADAR*), but also ADAR2 (*ADARB1*) which primarily edits at conserved sites in the genome (*51*, *53*, *54*). Using publicly available snRNAseq data from human brain, we confirmed the expression of *ADAR* in all major cell types including astrocytes (**Figure 5f)**. In astrocytes, the majority of expression was found in the astrocytic subtype expressing *VIM*, *SOX9* and *FOS*, with 19.4% of these cells expressing *ADAR*. The mean expression of *ADAR* also appeared to be higher in this subgroup, with *ADAR* among the top 500 most expressed genes. This is in keeping with *VIM*-positive astrocytes being immunoreactive, associating with response to toxins, viruses and cytokines, activation of surrounding neurons, projections over extended distances in the CNS, and astro-vascular interactions (*55–57*).

To explore whether the changes in A-to-I RNA editing seen in-vitro were also reflected in PD-affected human brain tissue, we explored RNA editing in a publicly available dataset of 5 control and 7 PD post-mortem brain samples (*58*). Using high depth RNA-seq data generated from the anterior cingulate cortex, and after controlling for covariates, we found that PD brain samples had significantly higher levels of RNA editing than control samples (beta 0.011, 95%, p <2.2×10^-16^ CI [0.009, 0.012]). This association remained significant in a further analysis, which also included the total number of editing sites per sample (beta 0.007, 95% CI [0.006, 0.009]). Next, we assessed the cell type specificity of the editing response. Focusing on genes with the greatest increase in mean editing rate in PD (top 50 genes), we used expression weighted cell type enrichment analysis (*59*) to formally assess the cell type-specificity of this gene set. We found significant enrichments for multiple glial cell types, including OPCs and oligodendrocytes. However, the cell type with the most significant enrichment was the astrocytes (3.8 standard deviations increased from the mean, adjusted p value 0.0014), suggesting that astrocytes are involved in this process in-vivo (**Figure 5g**) though not exclusively.

## Discussion

Astrocytes are the most abundant glial cells, supporting neuronal health and CNS immune responses through multiple heterogeneous reactive and proliferative states (*60*). Whilst several triggers for these states are well recognised, it remains less clear how different glial states contribute in the context of diseases associated with proteinopathies. Here we generated a platform of human iPSC-derived astrocytes to investigate the intersection between protein aggregation as a trigger for astrocytic state switching in disease. We used a serum-free small molecule approach to generate functionally active astrocytes, demonstrating homeostatic calcium responses, glutamate uptake, and maturation of neuronal function, as well as responses to inflammatory stimulation by canonical microglial triggers (LPS). Adopting single cell sequencing to characterise the molecular identity of the astrocytes revealed two astrocytic subclasses,protective and inflammatory,recapitulating key astrocytic states seen in the CNS and associated with PD (*13*). Furthermore, the transcriptomic signatures of our iPSC-derived astrocytes correlated highly with iPSC-derived astrocytes generated by other groups (*27*). Our cellular platform may therefore be used to model astrocytic states and sub-states, and uncover the mechanisms that underlie them, despite the inherent limitation of their developmental fetal phenotypes (*61*).

The innate immune response recognizes pathogen- and danger-associated molecular patterns (PAMPS & DAMPS) via pattern recognition receptors (including toll-like receptors (TLRs), and retinoic acid-inducible gene I (RIG-I)-like receptors) (*62–64*). The role of α-syn as a PAMP/DAMP is not well understood. It is currently known that α-syn may be transferred from neurons to astrocytes in vitro (*65*), or from astrocytes to neurons (*66*), and several mechanisms of astrocytic uptake and transfer have been proposed, including endocytosis (*67*) and tunnelling nanotubes (*68*), or via Toll like receptor 2 activation enhancing the uptake of fibrils (*69*). We previously demonstrated that physiological concentrations of oligomers provoke an immunological response that is largely TLR4 dependent, and that glial TLR4-Myd88 signalling in the substantia nigra may be causative in PD pathogenesis (*18*).

Here, we utilised the same α-syn aggregate species to reveal the downstream consequences of recognition by TLRs, and the underlying mechanisms of astrocytic reactivity driven by protein aggregation. Our data confirms that oligomers are recognised as DAMPS/PAMPs (driven by genes *TLR3, DDX58 (*RIG-I)). Typically, this is followed by activation of key inflammatory transcription factors, nuclear factor κB (NF-κB) and interferon regulatory factors (IRFs) that induce the release of type I interferon (IFN). The initial type I IFN release consequently induces the phosphorylation of IRF7, and phosphorylation of STAT2 and STAT1, which forms a complex with IRF9, known as the IFN-stimulated gene factor 3 (ISGF3). Oligomers induce the expression of *NFκB, IRF7, STAT1, STAT2, and ZBP1,* reflecting this pathway in the hiPSC astrocytes. Finally, the DAMPS/PAMPS trigger the activation of the NLRP3 inflammasome that facilitates the activation of caspase-1 and release of proinflammatory cytokines interleukin-1β (IL-1β), IL-18, and pyroptosis, a sequence also triggered by oligomers (*NLRP3, CASP1, IL1B)*. This transcriptomic response was mirrored functionally with the morphological switch to an activated astrocyte, release of inflammatory cytokines, and generation of reactive oxygen species. Moreover, in this reactive state, the astrocytic supportive function of promoting neuronal activity was lost, and neuronal toxicity was induced. Taken together these results suggest a mechanism whereby αsyn-O’s can trigger an inflammatory response in astrocytes with resulting glial and neuronal toxicity.

Intracellular double-stranded (dsRNA), evolutionarily a sign of viral infection, can also act as a DAMP, and dsRNA triggers multiple cytoplasmic receptors including ZBP1, MDA5, OAS, PKR which activate various arms of the innate immune cascade, including NF-κB signalling and the type 1 interferon response. Of the broad changes in gene expression observed in our experimental paradigm, oligomer induced type 1 interferon changes were accompanied by the activation of anti-viral response pathways, with upregulation of the cytosolic double-stranded nucleic acid (dsNA) sensing machinery. This machinery includes activation of the negative regulator ADAR1, an enzyme that performs RNA editing, breaking the homology of dsRNA, and thus dampening the immune response triggered by cytosolic dsNA and type 1 interferons (*45*). It is important in responding to cytosolic dsRNA that is endogenous in origin, and not secondary to viral infection. From an evolutionary perspective, this process is especially important in humans, where expansion of retrotransposition of non-coding repetitive elements,especially alu repeats, has markedly increased the amount of endogenous dsRNA within cells (*70*). The importance of the immune-dampening effect of ADAR-p150 is demonstrated in loss of function mutations which result in Aicardi Goutieres Syndrome, an infantile inflammatory encephalopathy which can also cause striatal necrosis, a brain structure implicated in PD (*71*).

ADAR1 has two major isoforms, with the p110 exclusively found in the nucleus and the p150 being largely cytoplasmic. In the hIPSC astrocytes, we observed promoter switching with the induction of the inflammatory isoform of ADAR1-p150. This resulted in a marked increase in A-to-I RNA editing site number and editing rate per site, most prominently in astrocyte-containing samples. The changes in editing were enriched in 3’UTRs suggesting that treatment with αsyn-O resulted in a higher proportion of editing activity within the cytoplasm, where ADAR1-p150 is known to localise. Furthermore, amongst genes that were both differentially edited and expressed, we found a significant enrichment for terms linked to viral infection and immune response in both the astrocyte-containing cultures. Thus the RNA editing is likely to dampen the inflammatory response to the oligomers in vitro.

Finally, we explored the role of RNA editing in PD. Using publicly available high depth RNA-seq data generated from the anterior cingulate cortex from control and PD-affected individuals, we found that PD brain samples had significantly higher levels of RNA editing than control samples. Having identified astrocyte specificity of the RNA editing response in vitro, we assessed the cell type-specificity of the editing response in vivo. Genes with the most significant editing in PD were enriched in multiple glial cell types, but most significantly in astrocytes, confirming the role of astrocyte RNA editing in vivo.

Our work raises a number of outstanding questions: how do the structural motifs of the oligomers act as danger associated molecular patterns, and trigger the interferon and editing response, and when does this process occur in the natural history of PD? However, most importantly, is A-to-I RNA editing a beneficial compensatory response to the inflammatory cascade, or does it exacerbate neurodegeneration in PD? Finally, whilst there is overlap between RNA editing and the genetic risk of PD (*72*), the role of altered RNA editing in PD remains unknown.

The findings here provide new insights into the mechanism by which inflammation may be implicated in PD pathogenesis: specific protein aggregates of α-syn may act as a DAMP to astrocytes, trigger inflammation and interferon like responses, which in turn triggers anti-viral dsRNA responses, leading to activation of RNA editing to dampen proteinopathy induced inflammatory responses. In this work, the disease specific trigger for this mechanism was the beta sheet rich, soluble oligomer of α-syn. However, such mechanisms may also be triggered by viral infections, which are believed to be associated with an increased risk of developing PD (*73*). The identification of dsNA sensing pathways provides a potential convergent mechanism between proteinopathy and viral infections in the pathogenesis of PD.

## Materials and methods

### Aggregation of human recombinant alpha-synuclein

Human recombinant α-Syn Monomeric WT or A53T α-Syn was purified from Escherichia coli as previously described (*74*). Aggregation reactions were carried out using a solution of α-Syn: 70 μM in 25 mM Tris buffer supplemented with 100 mM NaCl, pH 7.4 (in the presence of 0.01% NaN3 to prevent bacterial growth). The buffer was freshly prepared before each experiment and passed through a 0.02 μm syringe filter (Anotop, Whatman) to remove insoluble contaminants. Prior to incubation, the reaction mixture was ultra-centrifuged at 90k r.p.m. for 1h at 4°C to remove potential seeds. The supernatant was collected and separated in two fractions: one kept at 4°C at all times until use (monomers), and a second incubated in the dark at 37°C and 200 r.p.m., for ∼7–8 hours to generate oligomers, and avoid fibril formation. α-Syn was always kept in LoBind microcentrifuge tubes (Eppendorf, Hamburg, Germany) to limit surface adsorption.

### hiPSc culture

hiPSCs were derived from donors who had given signed informed consent for the derivation of hiPSC lines from skin biopsies as part of the EU IMI-funded program StemBANCC and reprogrammed as described (*75*). Briefly, the Cyto Tune-iPS reprogramming kit (Thermo Fisher Scientific) was used to reprogram fibroblasts through the expression of OCT4, SOX2, KLF4 and c-MYC by four separate Sendai viral vectors. Control 1 (C1) and 2 (C2) were derived by StemBANCC from an unaffected volunteer and control 3 (C3) was purchased from Thermo Fisher Scientific. Control 4 & 5 (C4 & C5) were purchased from Applied Stem Cell. hiPSCs were maintained on Geltrex in Essential 8 medium (Thermo Fisher Scientific) and passaged using 0.5mM EDTA.

### Differentiation of hiPSC into neurons and astrocytes

Differentiation of cortical region-specific astrocytes was performed using a modified protocol based on (*76*, *77*). Briefly, as demonstrated in Fig 1b, hiPSC were differentiated into neural precursor cells (NPCs) using an established protocol (*78*). In order to derive glial precursor cells (GPCs), NPCs were cultured with dual SMAD inhibition for 25-30 days, followed by culturing with the neural induction medium supplemented with 20 ng/ml human FGF-2).(*78*) The passage was performed twice per week (1:2 or 1:3) using Accutase (Cat #A1110501, Thermo Fisher Scientific) by vigorously breaking pellets to remove neuronal cells. Upon the appearance of glial morphology (around day 90 from the neural induction), the GPCs were cultured for 7 days with 10 ng/ml bone morphogenetic protein 4 (BMP4) and 20 ng/ml leukemia inhibitory factor (LIF) which activates the JAK/STAT signalling pathway, refreshing the medium every other day. On the 8th day, BMP4 and LIF were withdrawn, and the GPC were further differentiated for maturation in the neural induction medium without human FGF-2. 3-4 times more passages are required (1:3 or 1:4) until the complete loss of the precursor property, including proliferation.

For neurons, at around 35 days of induction, cells were dissociated into a single cell using accutase and approximately 150,000number of cells plated either PDL and laminin-coated glass bottom 8-well slide chambers (Ibidi/Thistle, cat No.80826), Geltrex coated 8-well ibidi chambers (cat No. IB-80826) or 96-well plates (Falcon, cat No. 353219). Medium was replaced every 4–5 days and cells were used at 60–90 days after induction.

### Live-cell imaging

Live-cell imaging was performed using an epi-fluorescence inverted microscope equipped with a CCD camera (Retiga; QImaging) or confocal microscope (ZeissLSM710 or 880 with an integrated metal detection system). For epi-fluorescence inverted microscope, excitation was provided by a xenon arc lamp with the beam passing through a monochromator (Cairn Research) and emission was reflected through a long-pass filter to a cooled CCD camera and digitized to 12-bit resolution (Digital Pixel Ltd, UK) and the data were analyzed using Andor iQ software (Belfast,UK). For confocal microscopes, illumination intensity was limited to 0.1-0.2% of laser output to prevent phototoxicity and the pinhole was set to allow optical slice at approximately 1–2μm. Pre-room temperature warmed HBSS was used as a recording buffer. 3–6 fields of view per well and at least 3 wells per group were used to analyze using ZEN, Volocity 6.3 cellular imaging or ImageJ software. All experiments were repeated at least 2-3 times with different inductions.

To measure Reactive Oxygen Species (ROS, mainly superoxide), cells were washed and loaded 2uM dihydroethidium (HEt, Thermo Fisher Scientific) in the recording buffer. The recording was performed using an epi-fluorescence inverted microscope equipped with 20x objective after a quick loading in order to limit the intracellular accumulation of oxidized product and the dye was present throughout the imaging. Excitation was set up to 530 nm and emission recorded above 560 nm was assigned to be for the oxidized form, while excitation at 380nm and emission collected from 405nm to 470nm were for the reduced form. The ratio of the fluorescence intensity resulting from its oxidized/reduced forms was quantified and the rate of ROS production was determined by dividing the gradient of the HEt ratio after the application of recombinant α-Syn against basal gradient.

For [Ca2+]c imaging, Fura-2, AM which is a ratiometric dye with a high affinity for Ca^2+^ was used. The cytosolic Ca2+as well as the rapid transient kinetics and decay times were assessed. 5u MFura-2 was loaded for 40 min and then washed twice before imaging. The fluorescence measurement was obtained on an epifluorescence inverted microscope equipped with a 20x objective. [Ca2+] was monitored in a single cell by obtaining the ratio between the excitation at 340nm (high Ca2+) and 380nm (low Ca2+) for which fluorescence light was reflected through a 515nm long pass filter. To trace morphological changes, Fluo4 was used and recorded using confocal microscopy.

Cell death was detected using SYTOX™ Green(SYTOX, Thermo Fisher Scientific) which is excluded from viable cells but exhibits red fluorescence following a loss of membrane integrity and Hoechst 33342 (Hoechst, Thermo Fisher Scientific) which stains chromatin blue in all cells to count the total number of cells. 500 nM SYTOX and 10 uM Hoechst were directly added into the dishes, and cells were incubated for 15 min. The fluorescent measurements were using confocal microscopy. Hoechst and PI were excited by 405nm with the emission between 405nm and 470nm. SYTOX was excited by a 488 nm laser with the emission between 488nm and 516nm. Percent cell death was quantified by the percent between the number of red fluorescent cells in the total number of Hoechst 33342 expressing cells per image.

### Immunocytochemistry

Cells were fixed in 4% paraformaldehyde and permeabilized with 0.2% Triton-X 100. 5% BSA was used to block non-specific binding before cells were incubated with primary antibodies either for 2 hours at room temperature or overnight at 4°C. The next day, cells were washed three times with PBS and incubated with a secondary antibody for 1hr at room temperature. Cells were mounted with an antifading medium after three times wash steps (DAPI was added in the second wash if required) and let dry overnight.

Lists of primary antibodies used; Anti-GFAP antibody (abcam, ab7260, 1:500), Anti-beta III Tubulin antibody (abcam, ab78078, 1:500). Lists of secondary antibodies used;Goat Anti-Chicken IgY H&L (Alexa Fluor® 488) (abcam,ab150169, 1:500), GoatAnti-Mouse IgG H&L (Alexa Fluor® 555) (abcam, ab150114, 1:500).

### Electrophysiology

Patch-clamp recordings of iPSC-derived neurons were performed using an infrared differential interference contrast (DIC) imaging system on an Olympus BX51WI upright microscope (Olympus, Japan) coupled with a Multipatch 700B amplifier under the control of pClamp 10.2 software package (Molecular Devices, USA), as described in detail previously.(*79*, *80*) For the recordings, a neuronal culture or co-culture was plated on glass coverslips, placed in a recording chamber mounted on the microscope stage and constantly perfused with a physiological buffer medium. The perfusion medium contained (in mM) 126 NaCl, 2.5 KCl, 2 MgSO_4_,2 CaCl_2_, 26 NaHCO_3_, 1.25 NaH_2_PO4, 10 D-glucose and was continuously bubbled with 95% O_2_ and 5% CO_2_ (pH 7.4) and maintained at 30-33°C. Whole-cell recordings were performed using glass pipettes with a resistance of 3.5-6 MΩ when filled with the intracellular solution. This solution contained (in mM): 126 K-gluconate, 4 KCl, 4 MgCl_2_, 2 BAPTA, 4 Mg-ATP, 0.4 Na-ATP (pH 7.2, osmolarity ∼295 mOsmol). In the whole-cell (immediately after membrane breakthrough), iPSC-derived neurons were recorded for the resting membrane potential (Vrest), membrane capacitance (Cm), the membrane time constant (τm), and input resistance (Rin). To induce neuronal firing, a series of sub- and supra-threshold rectangular current pulses were applied with a stepwise-increased stimulus intensity at the Vhold set at −60 mV to −75 mV. The second protocol tested was a slow-ramp current injection, ramped up with a 100–200 pA/s slope. The analysis of the AP waveform was performed for the first AP only to quantify the threshold value, the spike amplitude, overshoot, the spike width (duration at half-maximal amplitude), the rates of depolarisation and repolarisation phases as previously described (*81*).

### Isolation of single cells

#### Collecting cell pellets for bulk RNA-seq

To collect cell pellets, samples were trypsinised or scraped from the culture surface and placed in a 15ml conical tube. These tubes were centrifuged at 800g in a refrigerated centrifuge for 5 minutes, and the culture media decanted. The pellet was resuspended in 10ml chilled PBS per tube by pipetting, then centrifuged again using the above parameters before decanting the PBS. For bulk RNA sequencing, the cell pellets were frozen on dry ice and stored at -80.

#### Collecting cell pellets for single cell RNA-seq

Samples for single cell RNA sequencing followed the procedure above, though instead of freezing, the pellets were resuspended in 1ml PBS, and 100,000 cells were transferred to a 1.5ml falcon tube. These were centrifuged at 1000 rpm for 3 min at 4°C, before resuspending cells in 20ul chilled DPS. 180ul chilled 100% methanol was added dropwise to the cells while gently vortexing to prevent the cells from clumping, before fixing the cells on ice for 15 mins.

### Single-cell RNA-sequencing data generation and processing

Between 2400 to 4000 cells were loaded for each sample into a separate channel of a Chromium Chip G for use in the 10X Chromium Controller. The cells were partitioned into nanoliter scale Gel Beads in emulsions (GEMs) and lysed using the 10x Genomics Single Cell 3′ Chip V3.1 GEM, Library and Gel Bead Kit. cDNA synthesis and library construction were performed as per the manufacturer’s instructions. The RNA was reversed transcribed and amplified using 12 cycles of PCR. Libraries were prepared from 10 µl of the cDNA and 13 cycles of amplification. Each library was prepared using Single Index Kit T Set A and sequenced on the HiSeq4000 system (Illumina) using 100 bp paired-end run at a mean depth of 20-50 million reads per cell. Libraries were generated in independent runs for the different samples.

The reads were aligned to the human reference genome (Ensembl release 93, GRCh38) using Cell Ranger v3.0.2. The analysis was carried out using Seurat v3.0(*82*, *83*) following Seurat’s standard workflow. Cells expressing fewer than 200 genes were excluded from the subsequent analysis. In addition, we excluded cells with more than 3000 detected genes to remove suspected cell doublets or multiplets. Given that certain cell types, e.g. neurons, naturally express higher levels of mitochondrial genes, we applied a 10% cut-off for the percentage of mitochondrial genes expressed to filter out likely apoptotic cells. Using default parameters of Seurat, data for each sample were log normalised across cells and the 2000 most highly variable genes identified. Using the canonical correlation analysis (‘CCA’) (*83*) to identify anchors, we integrated the samples using Seurat v3 (*82*, *83*), followed by regression of the effect of cell cycle and scaling of the data. Dimensional reduction was performed using 50 PCs. We used Clustree v0.4.4 and Seurat’s plot functions to visualise the expression of astrocytic and neuronal marker genes across different cluster resolutions (0.05 - 0.5 in 0.05 increments and 0.5 - 1.0 in 0.1 increments) (*84*). A clustering resolution 0.25 was selected, as it was the lowest resolution that explained the heterogeneity in the samples. The differentially expressed genes between the clusters of interest were identified using Seurat’s FindMarkers() and the default ’Wilcox’ test.

### Bulk tissue RNA-sequencing data generation and processing

Libraries for sequencing were prepared using the Illumina TruSeq Stranded mRNA Library Prep kit by loading 50 ng of total RNA into the initial reaction; fragmentation and PCR steps were undertaken as per the manufacturer’s instructions. Final library concentrations were determined using Qubit 2.0 fluorometer and pooled to a normalized input library. Pools were sequenced using the Illumina NovaSeq 6000 Sequencing system to generate 150 bp paired-end reads with an average read depth of ∼137 million paired-end reads per sample.

We performed pre-alignment quality control using Fastp (v 0.20.0) with default settings, for adapter trimming, read filtering and base correction (*85*). Processed reads were aligned to the GRCh38 human reference genome using 2-pass STAR (v 2.7.0a), with gene annotations from Ensembl v93 (*86*, *87*). Parameters were set to match ENCODE options except, we only retained uniquely mapped reads and used STAR’s default of a minimum 3 bp overhang required for an annotated spliced alignment. Post-alignment quality metrics were generated using RSeQC (v2.6.4) and MultiQC (v1.8.dev0) (*88*, *89*). We found that an average of 90.4% reads were uniquely mapped.

The processed reads were also quantified with Salmon (v 0.14.1) using the mapping-based mode with a decoy-aware transcriptome based on GRCh38 and Ensembl v93 as the reference (*90*). Salmon’s options correcting for sequence, non-uniform coverage biases (including 5’ or 3’ bias) and GC bias in the data were enabled and the R package tximport used to transform Salmon transcript-level abundance estimates to gene-level values (*91*). Pipeline source code can be found in https://github.com/RHReynolds/RNAseqProcessing.

### Deconvolution

Cell type proportions in the bulk tissue RNA-sequencing samples were estimated using Scaden (v1.1.2) (*34*). Scaden trains on simulated bulk RNA-sequencing samples, generated from tissue-specific single cell data, and predicts cell type proportions in bulk tissue RNA-sequencing data. The training data was generated using the raw counts and cell types based on the clustering 0.25 from the single cell data. Thereby we created 2,000 artificial bulk tissue RNA-sequencing samples by randomly selecting 3,000 cells from the total of 8,132 cells. Prediction of the cell type proportions were made using the default parameters and the Scaden developers’ recommendations. Following deconvolution, significant differences in the cell type proportions between the αsyn-O treated and basal astrocytes and co-culture were investigated using a paired t-test and multiple comparison correction using Benjamini & Hochberg method, per cell culture. This was only applied to cell types with proportions >= 0.01, which included Astrocyte clusters 1 and 2 and Neuron clusters 1, 2 and 3.

### Differential gene expression analysis

Sources of variation in bulk tissue RNA-sequencing data were assessed by performing principal component analysis on the gene level expression filtered to include genes expressed in all samples of each cell culture and treatment.

We found that cell culture and cell-type proportions were significantly correlated with the first PC (**Supp fig. 3b**). The individual, age and sex correlated with PC2. Individual, RIN and astrocyte cluster 2 correlated with PC3, while culture, individual, age and neuron cluster 5 were significantly correlated with PC4. The treatment applied to the cell culture correlated with PC5. Accordingly, PC axes 2, 3 and 4 were included as covariates in the model for differential expression and splicing analyses of the bulk-tissue RNA-sequencing data. Bulk-tissue differential gene expression was examined using DESeq2 (v1.30.1) (*92*), including only genes expressed in all samples within a cell culture and treatment group, collapsing across individuals in a cell culture and treatment group; and controlling for covariates. A cut-off of FDR < 5% was used to consider a gene as significantly differentially expressed.

### Differential splicing analysis

Differential splicing analysis was conducted using Leafcutter (v0.2.9) (*93*). It detects changes in alternative splicing events by constructing clusters of introns that share splice sites and determining the difference in intron usage, measuring differential splicing in terms of the change in the percent spliced in (ΔPSI). Splice junctions outputted by STAR were filtered to remove those with length < 25 nucleotides and regions that overlapped the ENCODE blacklist regions (https://github.com/Boyle-Lab/Blacklist/tree/master/lists) (*94*). The junctions were annotated using junction_annot() from dasper (*95*), classifying them into the following categories, (based on if one end (acceptor or donor) or both ends match the boundary of a known exon) - annotated, novel acceptor, novel donor, novel combination, novel exon skip, unannotated and ambiguous gene (mapped to more than 1 gene). Those annotated as ambiguous were excluded from this analysis. Leafcutter was run to identify intron clusters by excluding introns of length greater than 1 Mb and those that were supported by < 30 junction reads across all the samples or < 0.1% of the total number of junction read counts for the entire cluster. Differentially spliced clusters were identified pairwise, in treated vs untreated astrocytes and co-culture samples, using leafcutter’s default parameters and controlling for covariates as identified by the gene level expression. 40,892 and 44,390 clusters (that lie in a single gene) were successfully tested for differential splicing in the astrocytes treated vs untreated and co-culture treated vs untreated respectively. An intron cluster and its overlapping gene were considered differentially spliced at FDR < 0.05 if the intron cluster contained at least one intron with an absolute delta percent spliced-in value (|ΔPSI|) ≥ 0.1. Functional enrichment analysis was performed using clusterProfiler (v3.18.1).(*96*) Gene ontology over-representation analyses were run and comparisons between genelists made using compareCluster(). We analysed differentially expressed genes at FDR < 5% and with at least >= 2 fold change in expression and differentially spliced genes at FDR < 5% and |ΔPSI| ≥ 0.1.

ADAR’s differentially spliced transcripts were linked to the protein isoforms by first identifying the transcripts the differentially spliced junctions overlapped with. The junctions overlapped with transcripts ADAR-201 and ADAR-202. The number of amino acids in these transcripts matched to P55265-5 (synonym p110) and P55265-1 (synonym p150) in Uniprot (*97*) respectively. We further ran multiple sequence alignment of the amino acid sequences from UniProt and Ensembl (obtained from the in silico translated mRNA is translated) for each of the isoforms, that was a match. ADAR’s differentially spliced junctions were visualised with ADAR’s protein coding transcript structures using ggtranscript (*98*).

### A-to-I editing

RNA editing analysis was undertaken with JACUSA2 v2.0.2 (https://github.com/dieterich-lab/JACUSA2), leveraging GNU parallel v20230722 (https://www.gnu.org/software/parallel/) (*99–101*). This utilises a drichilet multinomial distribution to ascertain whether transcripts are edited at a genomic site in two modes: in ‘detect’ mode it compares transcripts against a reference genome identifying editing in individual samples; in ‘differential’ mode it will compare two samples against each other, looking at sites that are differentially edited in one sample compared to another. Noting that transcripts with increased editing might not be successfully mapped during the alignment step, multi-sample 2-pass mapping with STAR v2.7.9a (https://code.google.com/archive/p/rna-star/) was re-run, allowing a more generous mismatch rate of 16 base pairs per 100.(*86*) This did not increase the rate of multimapping during alignment. PCR duplicates were identified by samtools v1.13 markdup (https://www.htslib.org) (*102*). A-to-I editing was assessed in properly paired, non-duplicate reads, with settings to exclude any potential editing sites near the start and end of reads, indel positions and splice sites, as well as sites within homopolymer runs of more than 7 bases. An editing site was considered significant if it had an absolute z score greater than 1.96 (|z| ≥ 1.96). Replicates for each sample group were input to JACUSA2 detect to output a list of editing sites for each of the six groups: astrocytes untreated, astrocytes treated, co-culture untreated, co-culture treated, neuron untreated and neuron treated. Differences in editing sites and rates were explored using R (v 4.2, www.r-project.org). JACUSA2 differential was used to ascertain those sites that were differentially edited in treated samples of each cell line, compared to untreated. Edits were annotated with Ensembl’s variant effect predictor (VEP, v93.5, https://www.ensembl.org/info/docs/tools/vep), filtering duplicate results by consequence, and biotypes of interest (*103*). Where results were not derived with Ensembl VEP, a manual annotation was undertaken using the Ensembl GTF file, deriving genic location and biotype. Editing sites were also annotated with repeat motifs downloaded from RepeatMasker v4.0.5 (http://www.repeatmasker.org/) (*104*). Functional enrichment of the differentially edited genes (FDR< 5%) that were also differentially expressed (FDR< 5% & at least 2 fold change in expression) was examined using clusterprofiler (v3.18.1) (*96*).

### Post-mortem brain editing analysis

Post-mortem human brain samples were sourced from publicly available data, including bulk and single-nuclear RNA sequencing data from 5 control and 7 PD anterior cingulate cortex samples from donors with Braak stage 5-6 disease (*105*). Single-nuclear gene expression was explored in python 3.9 (www.python.org) using the pl.dotplot function from scanpy 1.7.2 (*106*). As with the cellular models, bulk transcriptomic samples were passed through the editing pipeline including trimming with fastp, and alignment with STAR allowing 16 base mismatches per 100, and then identification of editing sites using JACUSA2 as above. Using R 4.2.0, sites were filtered to include those present in at least 2 samples per PD and control group, and presence in both groups. Sites with editing rate greater than 0 and less than 1 were input into a linear regression as follows: Editing_rate ∼ Disease_Group + Sex + RIN. Expression weighted celltype enrichment (EWCE v1.11.3, <u>nathanskene.github.io/EWCE/index.html</u>) analysis was undertaken using specificity matrices previously derived for this dataset from single nuclear RNA sequencing results (*59*, *105*). The ranked gene list input to EWCE was defined by the genes with the greatest increase in mean editing rate, relative to all the genes where editing was detected including genes with editing rate of 1.

## Acknowledgements

We would like to thank the donors for their fibroblast and brain tissue donation. We would also like to thank the Francis Crick Institute Flow Cytometry, Advanced Light Microscopy, Advanced Sequencing, and Bioinformatics and Biostatistics STPs for their help and equipment in conducting and analysing the flow cytometry, fluorescence microscopy, and single-cell RNA-seq experiments. This research was funded in whole or in part by Aligning Science Across Parkinson’s [ASAP-000509 and ASAP-000463] through the Michael J. Fox Foundation for Parkinson’s Research (MJFF). AZW was supported through the award of a Clinical Research Fellowship funded by the Wolfson Foundation and Eisai Ltd. SG was supported by Wellcome (100172/Z/12/2) and is currently an MRC Senior Clinical Fellow (MR/T008199/1). MR was supported by the UK Medical Research Council (MRC) through her award of Tenure-track Clinician Scientist Fellowship (MR/N008324/1). This work was funded in part by a grant from MJFF (Project Title: Reacting to alpha-synuclein: how astrocytes cause neuronal loss in Parkinson’s Disease; Grant ID: 18004).

## Data availability

The data that support the findings of this study are available at https://zenodo.org/records/10608268. Bulk-tissue and single-cell RNA-sequencing data will be accessible through the European Genome–phenome Archive. The RNA editing results are available at https://zenodo.org/records/10630845. Protocols used in this study can be found in the repository Protocols.io and the DOIs can be found in Supplementary Table 14.

## Code availability

Code used for the analyses of bulk-tissue and single cell RNA-sequencing data is available at https://github.com/karishdsa/ipscAstrNeurCocul. Code for the RNA editing analyses is available at https://github.com/aaronwagen/Astrocytes_editing.

## Ethics declaration

iPSCs lines were obtained from a number of different sources, commercially, from repositories (part of the EU IMI-funded program StemBANCC), or generated from fibroblasts from in-house skin biopsies taken under informed consent.

## Supplementary figures

**Supp Fig 1.**
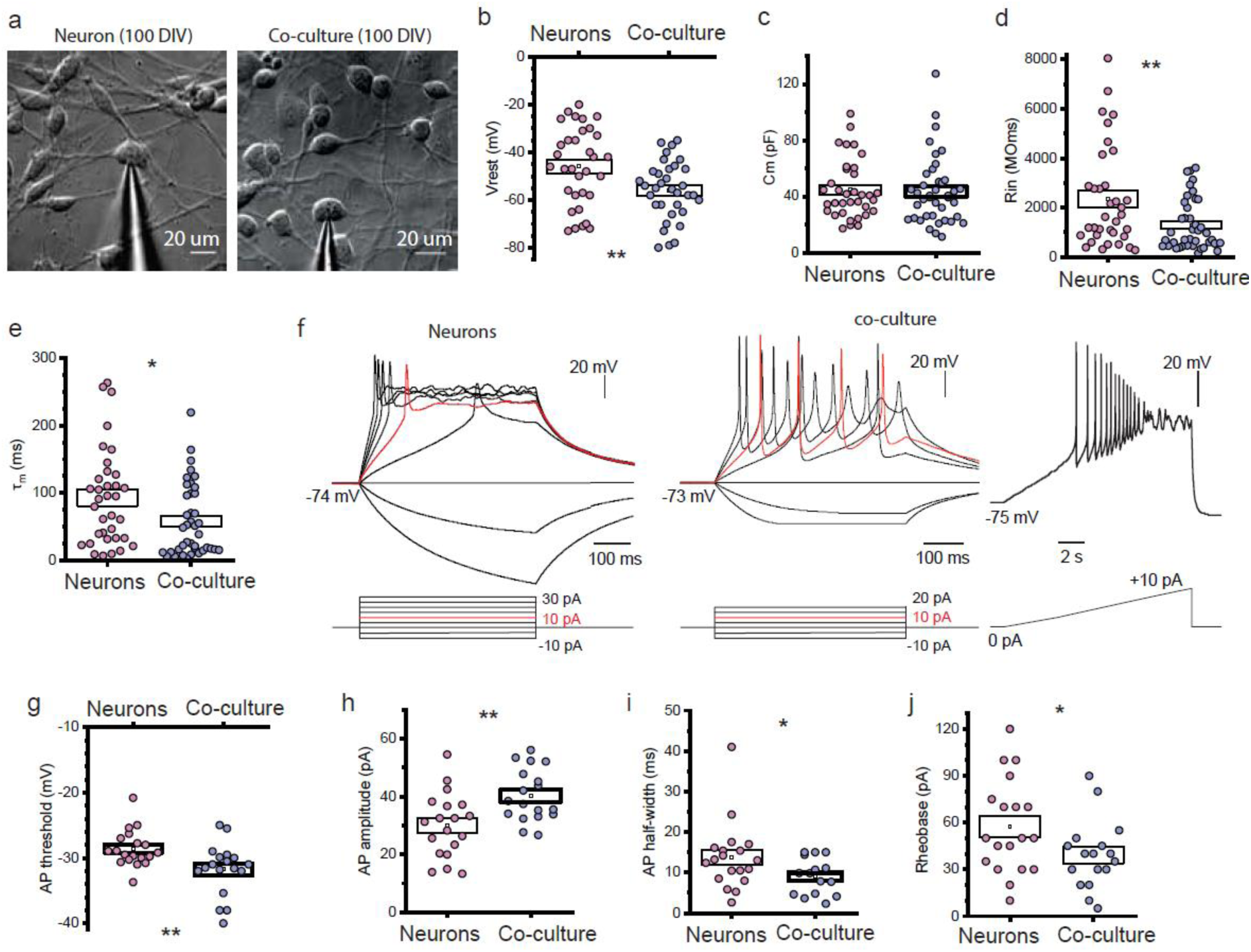
(a) Transmitted light images of iPSC-derived cells in neuronal culture (left) and neuron-astrocytic co-culture (right) for whole-cell recordings made from individual neurons using patch pipettes. (b) Statistical summary of the resting membrane potential (Vrest) revealed more hyperpolarised Vrest of iPSC-derived cortical neurons in co-cultures than in the corresponding age-matched cultures. (c - e) Passive membrane properties of iPSC-derived cortical neurons showed similar cell capacity (Cm, Fig 3.c), but different membrane constant (τm, Fig 3.e) and input resistance (Rin, Fig 3.d) between neuronal cultures and co-cultures, indicating a difference in biophysical maturation of the neurons. All data are mean with s.e.m. *P < 0.05, **p < 0.01 (the two-tailed unpaired t-test). (f) Representative recordings of action potential (AP) firing in an iPSC-derived neuron evoked by a series of rectangular depolarising current pulses of increased intensity (indicated on the bottom) in cultures. (Bii) Example recordings of AP discharge generated by an iPSC-derived neuron evoked by square depolarising current pulses (left traces) or a slow-ramp current (indicated on the bottom) in co-cultures. Note a train of evoked APs in co-cultures. Analysis of individual APs across iPSC-derived cortical neurons revealed significant changes in the threshold (g), spike amplitude (h), half-width kinetic parameters (i), and rheobase (j).

**Supp Fig 2.**
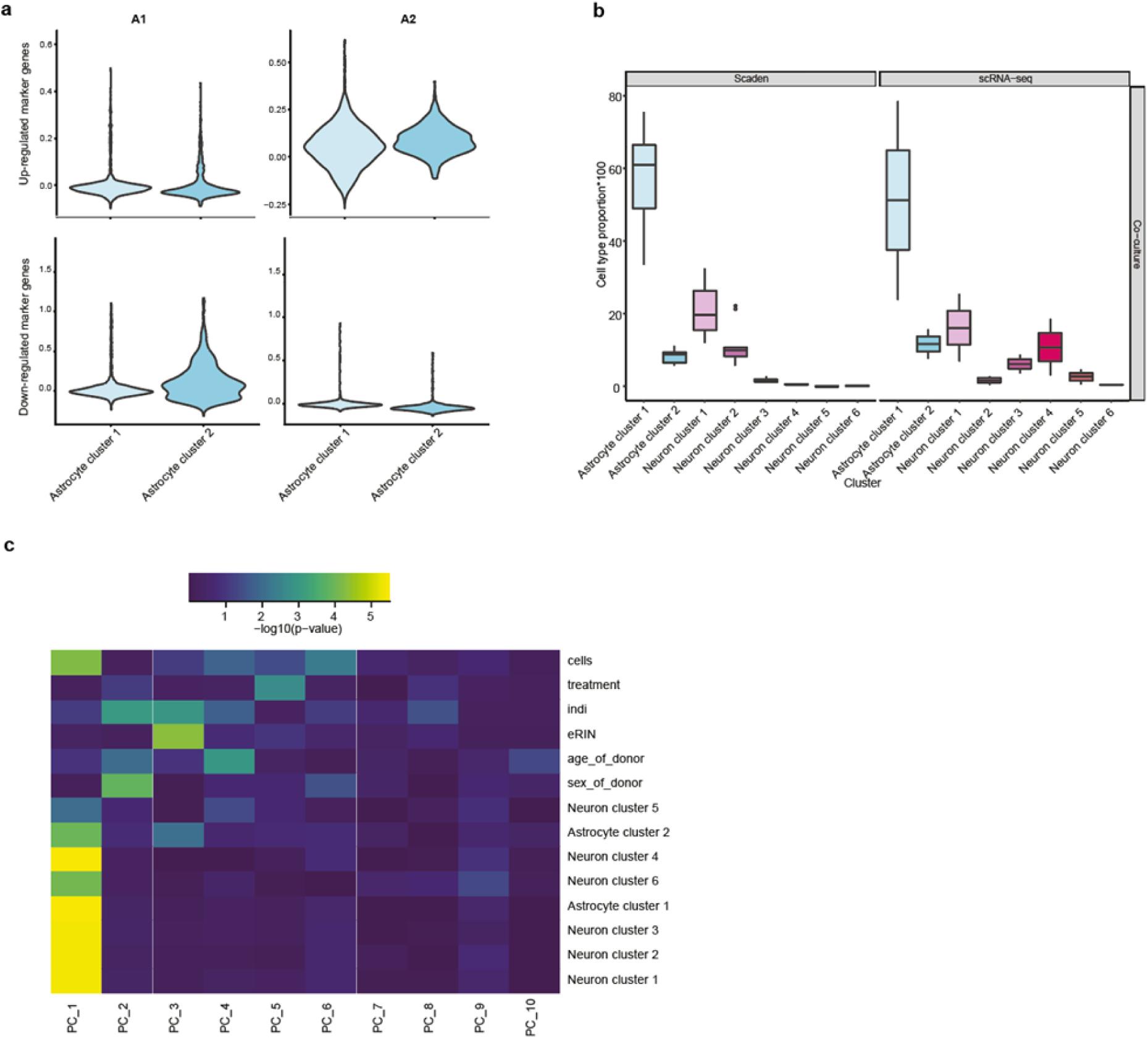
(a) Visualising the module score of the published up and down regulated A1 and A2 marker genes in astrocyte clusters 1 and 2. The module score, generated per cell for a set of genes using the AddModuleScore() in Seurat, is the difference between the average expression of each gene set and random control genes, per cell. (b) Cell type proportions estimated by the single cell data vs that predicted by Scaden’s deconvolution (c)Heatmap showing the correlation of the 1st 10 PCs with cell culture, treatment, individual, RIN, sex, age and the cell type proportions of the 8 cell types.

**Supp Fig 3.**
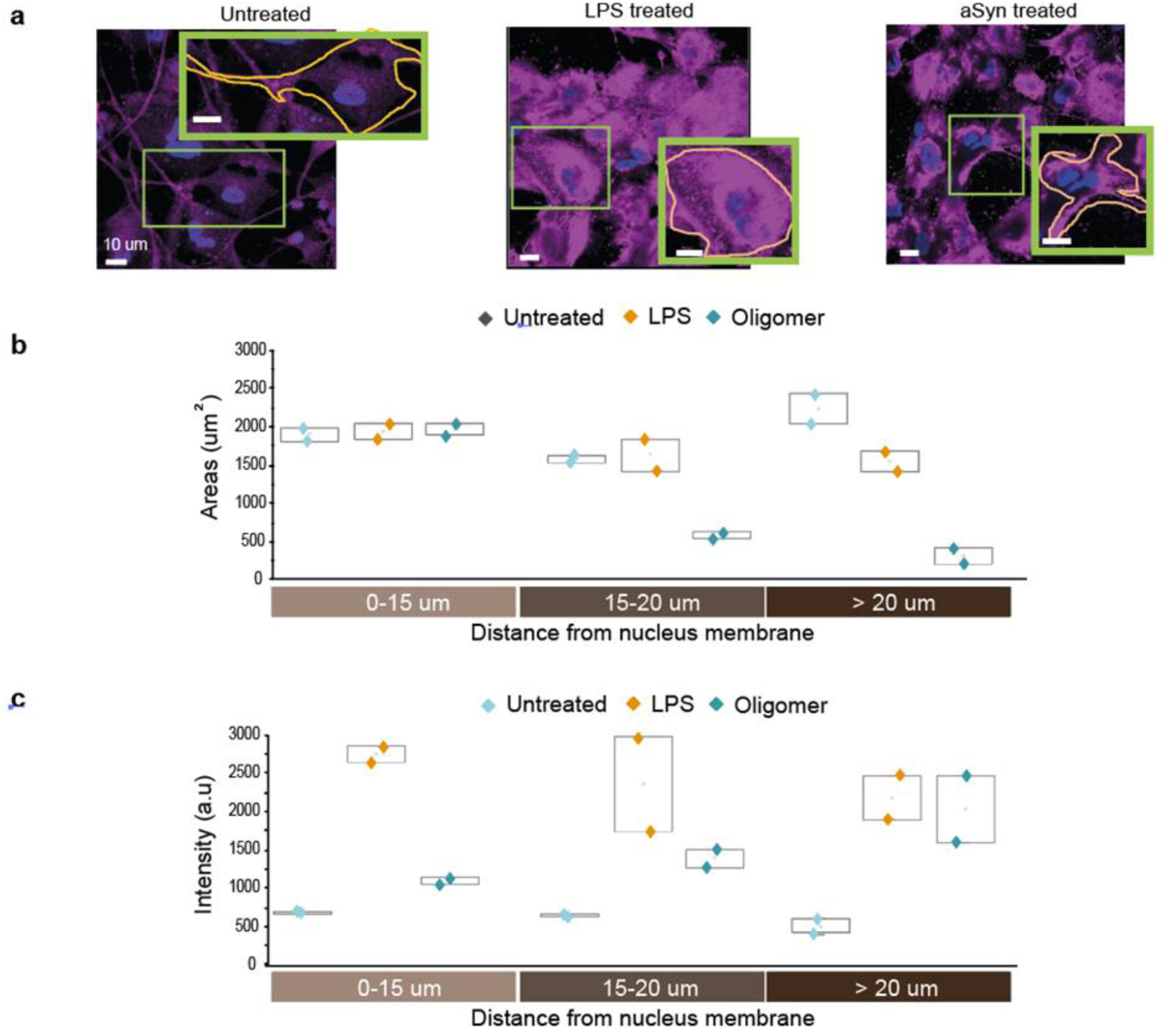
Morphological changes induced in reactive astrocytes on αsyn-O treatment: (a) representative images of astrocytes immune labelled with GFAP demonstrating change in morphology upon stimulation by LPS or αsyn-O. Assessment of morphology by segmentation of astrocytes followed by (b) assessing GFAP pixel area, according to distance from nuclear membrane (measurement of polarity) and (c) intensity according to distance from nuclear membrane.^38,39^

**Supp fig 4.**
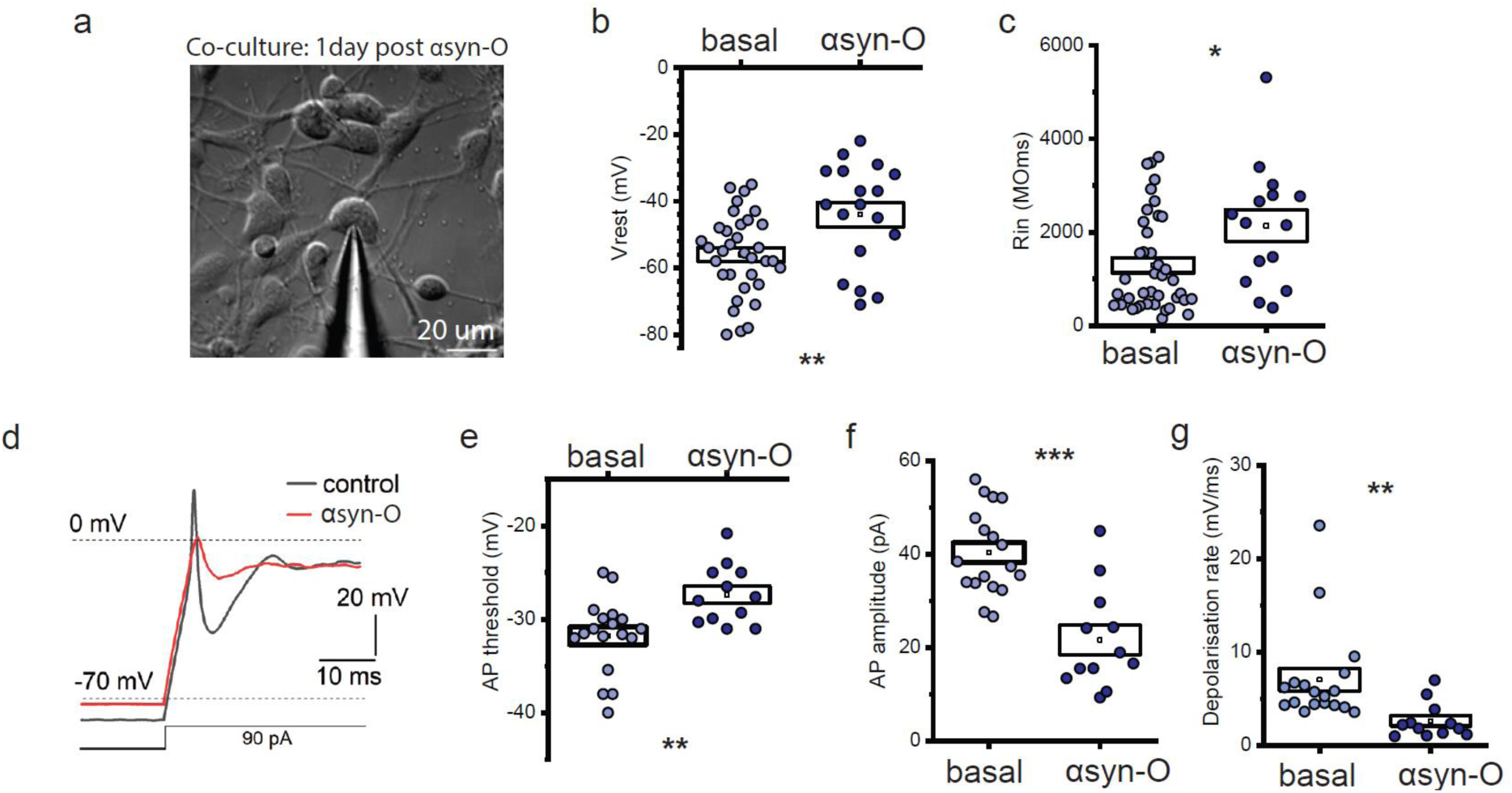
(a) DIC image of neuron-astrocytic co-cultures for whole-cell recordings of iPSC-derived neurons 1 day after treatment with exogenous mutant α-syn. (b) Analyses of membrane properties revealed a depolarised Vrest in iPSC-derived neurons at 1-day post-treatment with the pathogenic protein. (c) Cortical neurons exhibited an increased Rin (Input resistance) following the treatment compared with the parameters in control (untreated) co-cultures. (d) Representative recordings of action potential spike in a control co-culture (black) and 1 day post-treatment with the pathogenic α-syn (red). Pathological α-syn impaired the shape and kinetics of individual action potential spikes, as measured for the parameters of threshold (e), amplitude (f) and depolarisation rates (g).

**Supp Fig 5.**
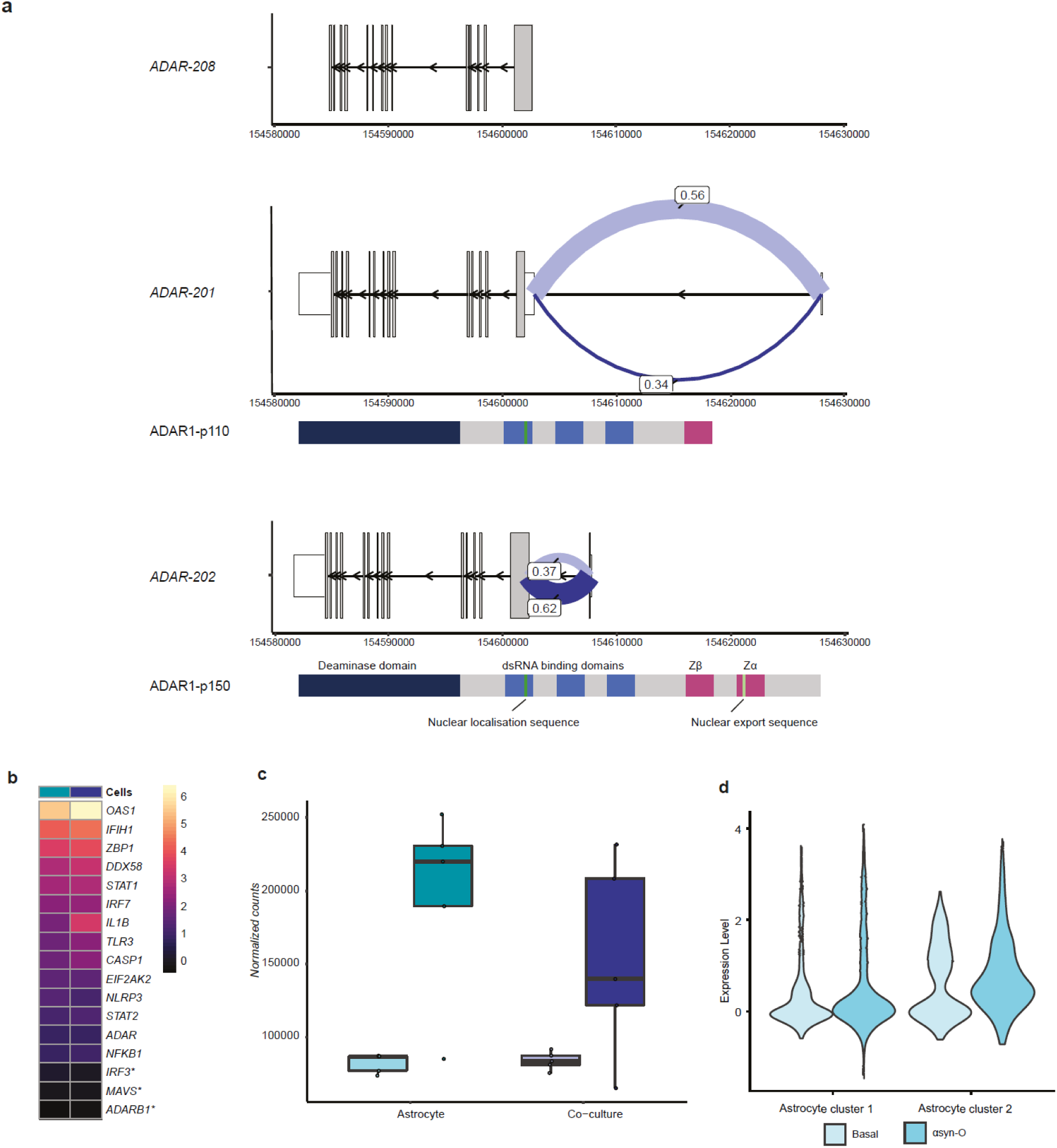
Differential gene expression and splicing of *ADAR* in co-culture on treatment with α-syn oligomers (a) Splicing in *ADAR* showing alternate isoforms. Junctions are labelled with the junction usage (b) Heatmap showing the log_2_FoldChange of the genes, in the innate immune response to dsRNA, (* not significantly differentially expressed at FDR < 5%) in the astrocytes and co-culture on αsyn-O treatment. (c) *ADAR* is differentially expressed in the astrocytes and co-culture on treatment with with α-syn oligomers (d) Expression of *ADAR* in the astrocyte cluster 1 and 2 in the single cell data

**Supp Fig 6.**
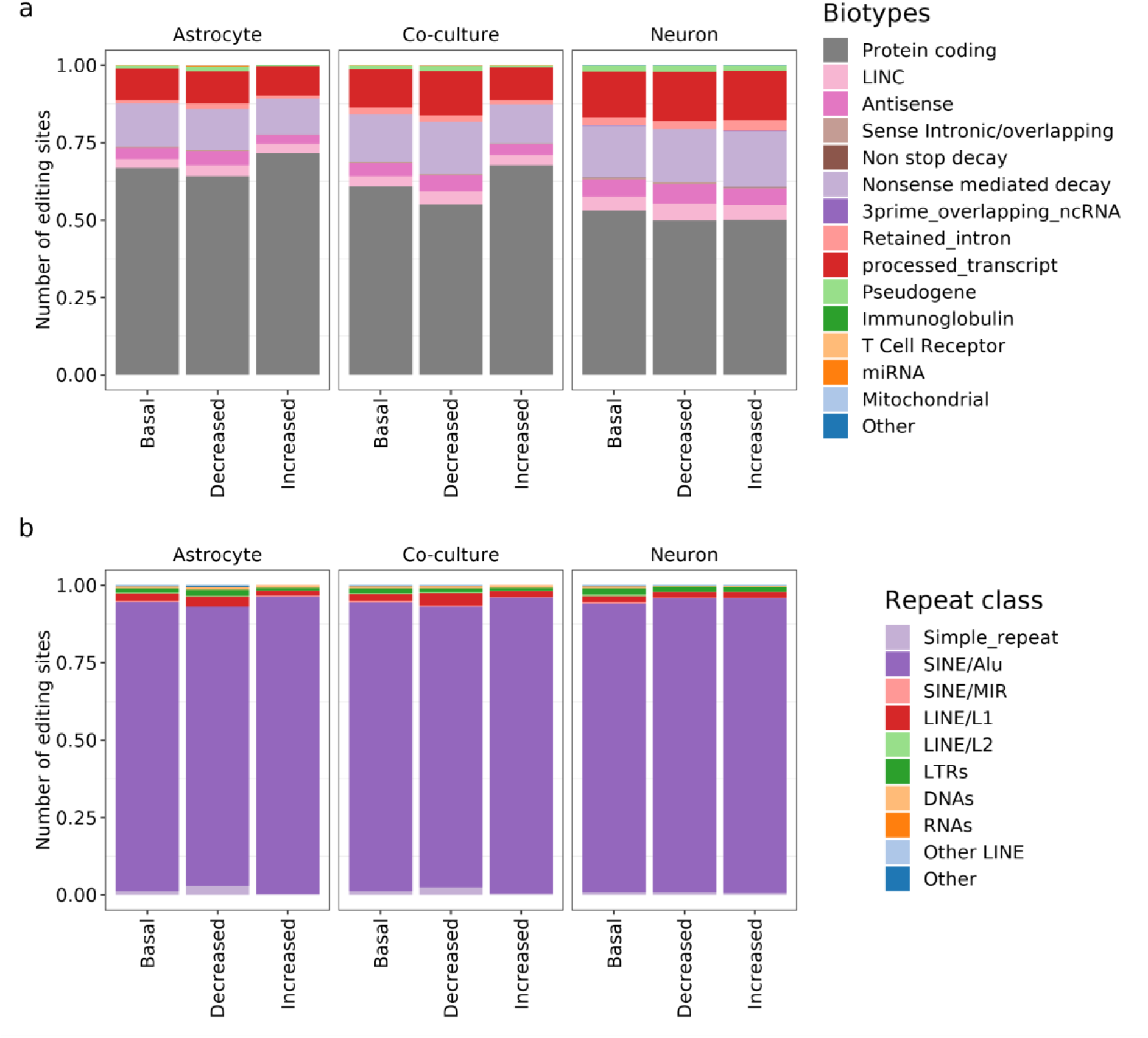
Changes in proportion of editing sites with α-synO treatment in astrocytes, co-cultures and neurons showing a) biotypes annotated with Ensembl Variant Effect Predictor (v93.5). b) Proportion of repeat regions annotated with RepeatMasker.

**Supp Fig 7.**
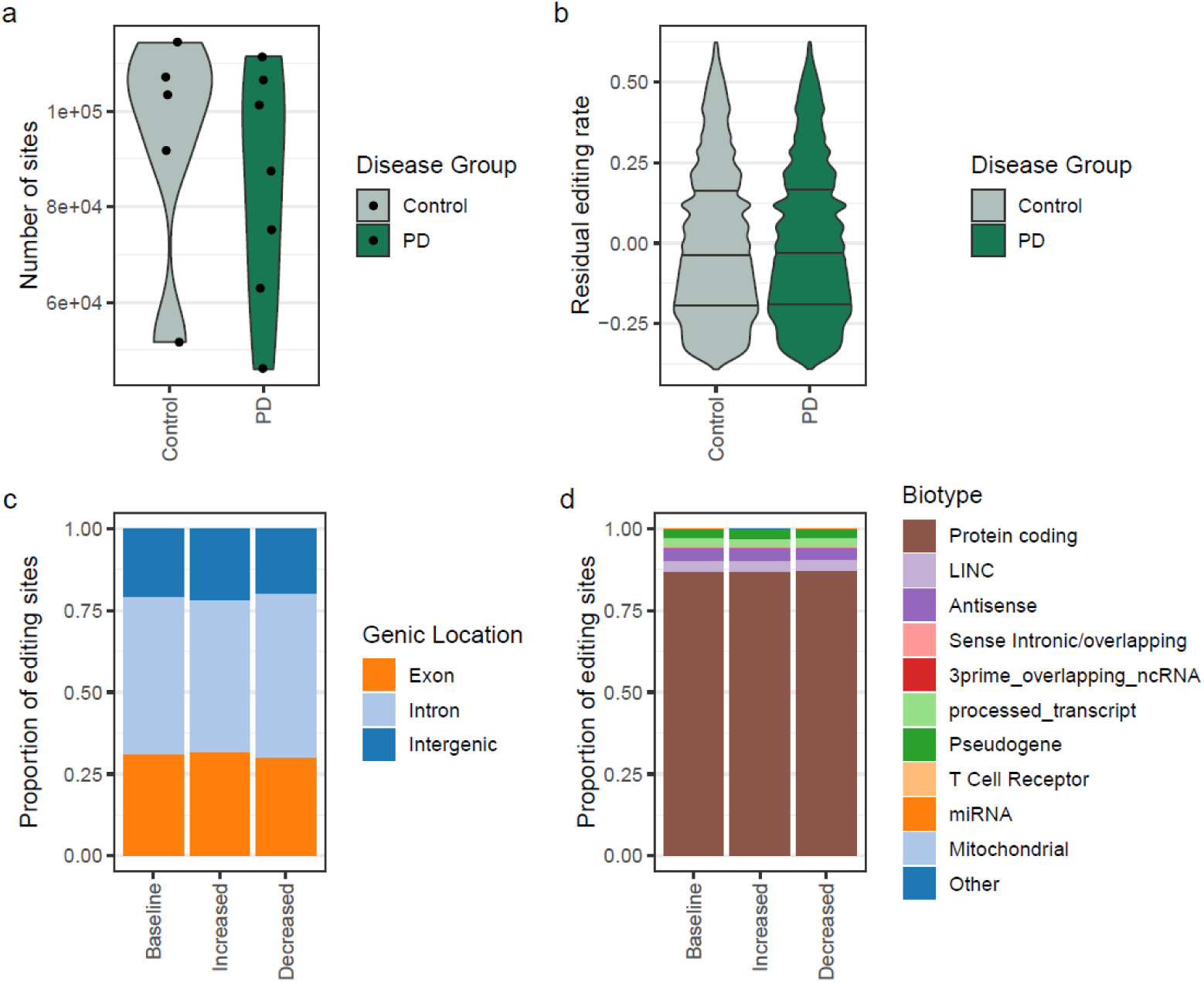
A-to-I RNA editing in PD affected human brain. (a) The total number of sites per brain sample. (b) Residual editing rate after correction for covariates. The change in the proportion of sites that have increased or decreased editing, relative to baseline, in genic location (c), and biotype (d).

